# Constructing a single-objective oblique plane microscope (OPM) for fast, multi-colour, high-resolution volumetric fluorescence imaging

**DOI:** 10.64898/2026.03.04.709686

**Authors:** Ziwei Zhang, Wenzhi Hong, Yunzhao Wu, Arpan Dey, Andrew Shevchuk, David Klenerman

## Abstract

Oblique plane microscopy (OPM) is a light sheet microscopy technique that uses a single high numerical aperture (NA) objective for both illuminating the sample and collecting emission fluorescence from a tilted plane within the specimen. OPM has become indispensable in biological and biomedical research, providing rapid, high-resolution volumetric fluorescence imaging of live cells and tissues while minimising phototoxicity and photobleaching. It also overcomes the sample mounting challenges associated with conventional light sheet microscopes that require two orthogonally placed objectives. However, the application of OPM has been limited by the complex design and the intricate optical alignment and characterisation needed, particularly with the remote-refocusing system (RFS) in the emission path. This protocol offers a detailed, step-by-step guide for constructing an OPM setup using commercially available components and for characterising its performance to ensure optimal imaging quality. We aim to deliver the unique merits of OPM to researchers in life science and medicine, enabling them to visualise the spatiotemporal organisation of key biomolecules, structures, and cells in 3D at high resolutions.

## Introduction

Fluorescence microscopy has revolutionised modern biology by enabling the direct visualisation of biomolecules and structures in cells or tissue, offering critical insights into diverse physiological processes, such as gene expression, protein interaction and transport, and cell division [1]. Most fluorescence microscopy techniques share a similar core principle: the molecule or structure of interest in the specimen is labelled with a fluorescent dye, which can be excited by light of certain wavelengths to emit fluorescence photons at a longer wavelength. The fluorescence is then collected, filtered, and recorded by a digital camera, producing an image of the labelled targets with high specificity and contrast.

Typical wide-field fluorescence microscopy illuminates the entire specimen and acquires two-dimensional (2D) images [2]. While straightforward, this illumination configuration presents two limitations: firstly, with all fluorophores in the specimen excited, the fluorescence from fluorophores outside the focal volume is also collected, resulting in a blurry background in the image [3]; secondly, prolonged light exposure can produce radicals (e.g., reactive oxygen species), which can damage the fluorescent molecules and live cells, ultimately limiting the duration and quality of live-cell imaging [4].

Light-sheet fluorescence microscopy (LSFM) has been developed to address the above challenges [5], [6]. Instead of illuminating the entire specimen, LSFM employs a thin sheet of light to illuminate the sample and only excites the fluorophores within this sheet, effectively eliminating the background from out-of-focus fluorescence and significantly enhancing the signal-to-noise ratio (SNR). More importantly, this optical sectioning capability allows for volumetric imaging by sequentially scanning the light sheet through the specimen. The resulting images obtained from different planes can be computationally stacked, enabling the reconstruction of three-dimensional (3D) structures in the specimen, thereby revealing spatial biological features that 2D imaging cannot capture. By using a sufficiently thin light sheet, LSFM may offer a high spatial resolution along the scanning direction. The plane-scanning method in LSFM also enables faster imaging with higher temporal resolution than the voxel-scanning approach used in confocal microscopy [7], allowing for the observation of dynamic biological processes in real time. Furthermore, the selective illumination of one plane at a time minimises photobleaching and phototoxicity, making LSFM ideal for prolonged live imaging of sensitive specimens, such as live cells, organoids, and embryos.

A conventional LSFM implementation employs two orthogonally aligned objective lenses [8]. One objective lens introduces a thin, horizontal light sheet to illuminate the specimen, while the other, positioned perpendicular to the illuminated plane, collects the emitted fluorescence. This configuration often requires specially designed sample holders, such as gel cylinders, to position the specimen [9]. Customised microscope body is also required to mount the two objective lenses. The inverted selective-plane illumination microscopy (iSPIM) [10] configuration obliquely illuminates the specimen from the top, eliminating the need for gel cylinders; however, this design can lead to spatial hindrance between the objectives, especially when the detection objective has a high numerical aperture (NA) and correspondingly short working distance (Figure 1B). This configuration also requires custom sample holders and is incompatible with standard containers in biological research, such as Petri dishes or well plates. Open-top light sheet (OTLS) is an inverted version of iSPIM and illuminates the sample from below through the coverslip (Figure 1C) [11]. OTLS bypasses the need for custom sample holders but still faces spatial constraints between the objectives. A shared limitation across these configurations is the requirement to physically translate the specimen, or the objectives, along the detection axis during volumetric imaging. This mechanical scanning process limits the acquisition speed and potentially introduces mechanical vibration and drift, resulting in reduced image quality and temporal resolution.

**Figure 1:**
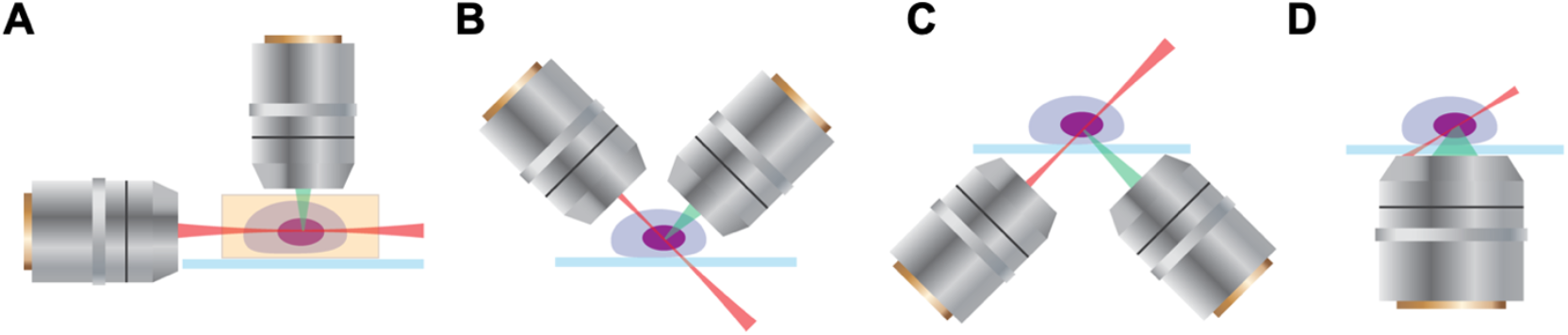
Configurations of different light-sheet fluorescence microscopy (LSFM) setups. (A) A traditional LSFM setup uses two orthogonally aligned objectives for illumination and fluorescence collection, respectively. (B) Inverted selective-plane illumination microscopy (iSPIM) illuminates the sample and collects the fluorescence using two objectives above the sample stage. (C) Open-top light sheet (OTLS) utilises two objectives below the sample for illumination and fluorescence collection. (D) Oblique plane microscopy (OPM) uses the same objective lens under the sample for sample illumination and fluorescence collection.

Oblique plane microscopy (OPM) was first developed in 2008 to overcome the limitations of conventional LSFM configurations (Figure 1D) [12]. OPM utilises a single objective lens for sample illumination and fluorescence collection, along with a remote-refocusing system (RFS) in the detection path for image restoration and acquisition. The sample is illuminated by an oblique light sheet from the primary objective lens. The fluorescence collected by the same objective lens passes through a secondary objective lens, creating an undistorted replica of the image plane at its focal plane. This replica is then imaged by a tertiary objective lens mounted at an oblique angle, such that the tilted image is corrected at the camera. By virtue of this design, OPM is compatible with most standard sample containers and commercialised microscope platforms, as well as high-NA primary objectives. OPM allows a relatively large imaging volume to be imaged at a high temporal resolution, ideal for live imaging (e.g., scan speed limited by the camera frame rate) at a high spatial resolution (limited by optical diffraction).

Following the original development of OPM, several advancements have further enhanced its imaging performance. The epi-illumination SPIM (eSPIM) improved the overall detection NA from ~0.7 in the original OPM setup to ~1.06 by introducing a water chamber between the secondary and tertiary objectives [13]. Another key advantage of eSPIM is the absence of mechanical sample translation during imaging, as the only moving component is a scanning galvanometric (Galvo) mirror that directs the light sheet through the specimen, while the sample stage and objective lenses remain stationary. The eSPIM setup was further improved by replacing the tertiary objective with a bespoke glass-tipped objective (AMS-AGY v1.0), resulting in no appreciable loss in NA [14]. In addition, by employing a high-NA oil-immersion primary objective lens and a Bessel beam, this design achieved higher spatial resolution in all three dimensions compared to the original eSPIM.

The eSPIM configuration and its variants provide a broad range of imaging volumes, spatial resolutions, and volume acquisition rates, supporting various biological applications. The original eSPIM achieved 3D imaging of live mammalian cells in multi-well plates [13]. Subsequent modifications expanded its capabilities to mesoscopic volumetric imaging and demonstrated whole-body imaging of live zebrafish larva blood flow at 5 Hz volume rate. DaXi, another variant of eSPIM, achieved a large imaging volume of 3,000×800×300 µm^3^ with a lateral resolution of 450 nm and an axial resolution of 2 µm [15]. Other eSPIM adaptations have enabled visualisation of diverse biological samples, including the brain of *Danionella cerebrum* [16], freely moving *Nematostella vectensis*, and *Drosophila* embryos [17], as well as subcellular components, such as actin cytoskeleton [18] and endoplasmic reticulum [19].

Despite the unique advantages of OPM and its variants, this technology remains inaccessible to most biology laboratories due to the lack of a commercialised product. Meanwhile, building an OPM-based microscope from scratch can be highly challenging. Firstly, selecting optical components (e.g., objective lens) that achieve optimal performance for certain applications requires previous experience with OPM construction. Secondly, critical expertise in optical design and alignment is required to practically build an OPM setup, since imperfections in alignment can accumulate along the optical path and lead to distortion or aberrations in the final image. Therefore, sophisticated validation and characterisation are essential throughout the alignment process to ensure optimal imaging performance. Thirdly, skills in hardware control and computational image reconstruction and processing are also required to operate the microscope and acquire images. Unfortunately, no detailed, step-by-step instruction on OPM construction and characterisation is available.

In this Protocol, we provide a comprehensive guide for constructing a compact single-objective OPM setup, with particular user-friendly instructions on the RFS, using commercially available optical components and digital devices. We will include detailed illustrations for alignment and characterisation to ensure robust performance and high imaging quality. This Protocol aims to break the technical barriers in assembling and utilising OPM and to enable researchers to conduct fast, high-resolution, and multi-colour volumetric fluorescence imaging of biological samples. Additionally, the principles and techniques outlined here may offer valuable insights for other LSFM methods incorporating RFS modules.

### Illumination methods in LSFM

Similar to many other fluorescence microscopy techniques, LSFM relies on two fundamental components: the illumination system and the detection system. Different combinations of these components lead to the innovation of various imaging modalities that fulfil specific applications. In LSFM, the illumination strategy is of critical importance as the choice of the light-sheet directly influences fundamental imaging quality, including image contrast, axial resolution, and optical sectioning capability [20]. The light-sheet generation methods differ in their underlying physical principles, producing sheets with distinct characteristics such as intensity profile, depth of focus, and interaction with different sample types. Each method represents a different solution to the fundamental trade-offs between these parameters, necessitating careful consideration based on experimental requirements. Here, we provide a thorough discussion on major light-sheet illumination methods, aiming to offer guidance on the choice of these methods for different applications.

#### Gaussian Light Sheets

The Gaussian light sheet represents the most fundamental and widely implemented illumination approach in LSFM. Gaussian light sheets are commonly generated by passing a collimated Gaussian beam through a cylindrical lens [21], [22]. The resulting illumination sheet exhibits typical Gaussian beam characteristics: the thickness of the sheet is the thinnest at its waist and gradually increases with propagation distance due to diffraction, while its intensity decays exponentially from the central axis to the periphery.

The advantages of Gaussian light sheets lie in their straightforward implementation and optimal optical sectioning capabilities. Their confined illumination minimises photobleaching and phototoxicity while enhancing image contrast by reducing out-of-focus background signals. Gaussian light sheets also improve axial resolution compared to epi-fluorescence illumination, as thinner light sheets can be generated by employing high-NA illumination objectives [20]. When paired with high-NA detection optics, this configuration further enhances light collection efficiency and lateral resolution [23].

However, Gaussian light sheets face an inherent trade-off between sheet thickness and propagation length. The useful illumination region is typically constrained to within the Rayleigh length, beyond which the beam rapidly diverges, compromising both optical sectioning and axial resolution [3]. Producing a thinner sheet to achieve superior axial resolution inevitably limits the imaging field of view (FOV) [24]. Additionally, high-NA objectives, while beneficial for producing thinner sheets, are limited by their short working distances, which impose constraints on the setup and sample mounting formats [25].

To address these limitations, thicker light sheets are often employed for applications requiring a larger FOV. Pairing with long-working-distance, low-NA objectives can also relax the constraints on detection optics but sacrifice resolution [26]. Balancing high resolution with an expanded detection FOV remains a significant challenge in LSFM [27]. In addition, due to the diffractive nature of Gaussian beams, the excitation light sheet degrades as it propagates through highly scattering samples, thereby limiting the penetration depth [28], [29]. As a result, Gaussian light sheets are mainly advantageous for live cell imaging and for studying thin and light-sensitive samples [30].

#### Scanned Bessel Beams

Bessel beams are a unique type of light-sheet illumination, characterised by their ability to maintain beam shape over extended distances [31]. They can be generated through methods such as passing a Gaussian beam through an axicon (a conical lens) [32], [33], an annular aperture [34], [35], [36], a spatial light modulator with the appropriate phase pattern [37], [38] or holographic optical elements [39], [40], resulting in an intensity profile consisting of a bright central lobe surrounded by concentric rings (side lobes). In most LSFM implementations, Bessel beams are scanned across the FOV to create a uniform illumination sheet. Two fundamental properties make Bessel sheets particularly valuable for biological imaging. First, their non-diffracting nature allows the central lobe to maintain its narrow profile over extended distances without significant divergence [41]. Second, they exhibit self-reconstruction capability, which enables the beam to reform after encountering obstacles in the light path [42]. These characteristics make Bessel beams especially effective for imaging thick biological samples or penetrating highly scattering media [43]. Research has shown that by carefully tuning the thickness of the Bessel beam’s central lobe to match the DOF of the detection objective, isotropic resolution of approximately 300 nm can be achieved over a 40 μm FOV along the beam’s propagation direction [35] - a performance unattainable with conventional Gaussian light sheets.

However, implementing Bessel sheets presents specific challenges. While the thin central lobe can provide high axial resolution, the inherent side lobes generate out-of-focus fluorescence that compromises optical sectioning and reduces image contrast. Mitigating these effects typically requires computational post-processing through deconvolution algorithms [44], adding complexity to the imaging analysis pipeline, or additional measures such as combining with structured illumination[35] or confocal detection [45]. In addition, the implementation of scanning Bessel beams demands more sophisticated optical alignment than conventional Gaussian sheets. While static line Bessel sheets have been developed, they retain greater technical complexity [46]. Despite these challenges, Bessel sheets have proven invaluable for applications requiring maintained resolution over extended FOV, particularly in deep tissue imaging.

#### Lattice Light Sheets

The lattice light sheet represents an advanced implementation of Bessel beam principles. Lattice light sheets are generated by the coherent superposition of multiple Bessel beams arranged in a periodic, grid-like pattern [47]. The lattice structure is typically created using a diffraction grating [48] or a spatial light modulator [47], [49]. Importantly, by using a linear array of Bessel beams with the correct periodicity, destructive interference occurs and suppresses the side lobes of individual Bessel beams, forming an optical lattice that has a more uniform and evenly distributed illumination profile.

A key strength of lattice light sheets lies in their ability to deliver ultrathin light sheets, often with an FWHM of ~1 μm, across a relatively large FOV of ~80 × 80 μm [35], [50]. The high optical efficiency of lattice beams coupled with high-NA detection objectives (up to NA = 1.1) not only improves resolution but also enables the use of faster cameras, such as scientific complementary metal-oxide semiconductor (sCMOS) cameras, thus enhancing the temporal resolution [27].

Lattice light-sheet microscopy (LLSM) operates in two distinct modes: dithered [47], [51] and structured illumination microscopy (SR-SIM) [52]. The dithered mode rapidly scans the lattice pattern to create uniform illumination, prioritising speed and sample viability. With an impressive imaging speed of up to 100-200 frames per second, this mode is particularly well-suited for capturing highly dynamic processes in living samples. The SR-SIM mode achieves superior resolution beyond the diffraction limit by collecting multiple images per plane, though at slower acquisition rates. While this mode offers better resolution than the dithered approach [53], its increased exposure times and light doses make it less suitable for extended live imaging, though it remains significantly faster than conventional scanning Bessel beam methods [54].

Despite these capabilities, LLSM faces certain limitations. The requirement for precise lattice pattern projection makes the system particularly sensitive to light-sheet degradation, limiting its effectiveness in penetrating deeper or more opaque tissues. The optical complexity and cost of LLSM systems also present barriers to widespread adoption. Simplified variants using physical apertures instead of spatial light modulators have emerged to reduce cost and complexity [46], though they may lack the side lobe suppression achieved with lattice periodicity [27][27], [55]. Nevertheless, LLSM is particularly suitable for long-term imaging of live samples, where high spatiotemporal resolution and minimal phototoxicity are essential. The availability of commercial, user-friendly systems has further expanded its accessibility to researchers without extensive optical expertise.

#### Airy Beams

Airy beams are commonly generated by applying a cubic phase mask to a Gaussian beam using a spatial light modulator [55]. Alternatively, an approximate Airy beam can be produced by precisely tilting cylindrical lenses [56]. Airy beams exhibit a characteristic banana-shaped trajectory and possess self-reconstruction properties similar to Bessel beams [57]. Their elongated profile enables uniform illumination over a large field of view, extending up to several hundred micrometres [58], making them well-suited for imaging large tissues [59]. In addition, the power of an Airy sheet is distributed over a larger area compared to an equivalent Gaussian light sheet, resulting in reduced peak intensity and, consequently, lower photobleaching [60].

One notable feature of Airy beams is their asymmetric excitation pattern. When the main lobe of the beam curves within the detection focal plane and its side lobes are symmetrically distributed on either side, the main lobe illuminates a flat plane but results in a thicker overall sheet [20]. Alternatively, when the beam is rotated 45° around the propagation axis, the illumination plane becomes curved, but some side lobes align to overlap with the volume illuminated by the main lobe, improving optical sectioning [55]. To accommodate the curved illumination volume, a detection objective with an extended depth of field (DOF) is favoured. This is typically achieved using objectives with a lower numerical aperture (NA), which consequently compromises both lateral and axial resolution [20].

Mitigating the effects of side lobes requires post-processing techniques such as deconvolution [27]. Other optical approaches, such as two-photon excitation [61], have also been developed however at the expense of higher complexity. For LSFM implementations, beam scanning is required to generate a uniform Airy sheet, posing complexity on the alignment. Despite these challenges, Airy beams provide an effective balance between a large FOV, high optical sectioning, and contrast, making them highly advantageous for imaging large, live samples, particularly when minimising phototoxicity is critical.

The choice of light-sheet type depends on the balance between resolution, phototoxicity, field of view, and sample thickness, and it should be tailored to specific research requirements. While in LSFM, the lateral resolution is entirely dependent on the NA of the detection objective, the axial resolution is governed by the product of the axial excitation intensity profile and the detection PSF [60]. Gaussian light sheets exhibit anisotropic resolution. High axial resolution can be obtained by imaging with a thin sheet and is inversely related to the sheet length. However, for conventional Gaussian light sheets, the thickness of the sheet may exceed the DOF of the detection PSF. In this scenario, axial resolution is mainly dominated by the detection NA, and the thicker sheet contributes to the out-of-focus background. In contrast, Bessel and lattice configurations can achieve and maintain isotropic or near-isotropic axial resolution over longer distances due to their thin main lobe thickness [35], [62]. Airy beams provide good axial resolution but typically require computational reconstruction to achieve optimal results.

The effective optical sectioning also varies: Gaussian sheets offer the best sectioning for simpler implementations, while multi-lobe beams like Bessel and lattice configurations have inferior sectioning compared to their axial resolution because they illuminate out-of-focus planes, leading to a background that blurs the resulting image.

From a practical perspective, the complexity of light sheet generation increases significantly when transitioning from simple Gaussian sheets to advanced techniques like lattice light sheets. This progression demands higher precision in optical alignment, enhanced system stability, and incurs greater costs, especially when incorporating spatial light modulators and sophisticated control systems.

In summary, Gaussian light sheets are well-suited for imaging cellular samples with a focus on optimal contrast and sectioning, while the self-reconstructing properties of Bessel and Airy beams make them advantageous in imaging thick, highly scattered tissues. Lattice light sheets provide uniform illumination, making them ideal for live-cell imaging and dynamic studies, though they require more advanced and expensive equipment. A comprehensive comparison of these methods is presented in **Table 1**; their applications in investigating various biological samples and systems are summarised in **Supplementary Table S1**.

**Table 1:**
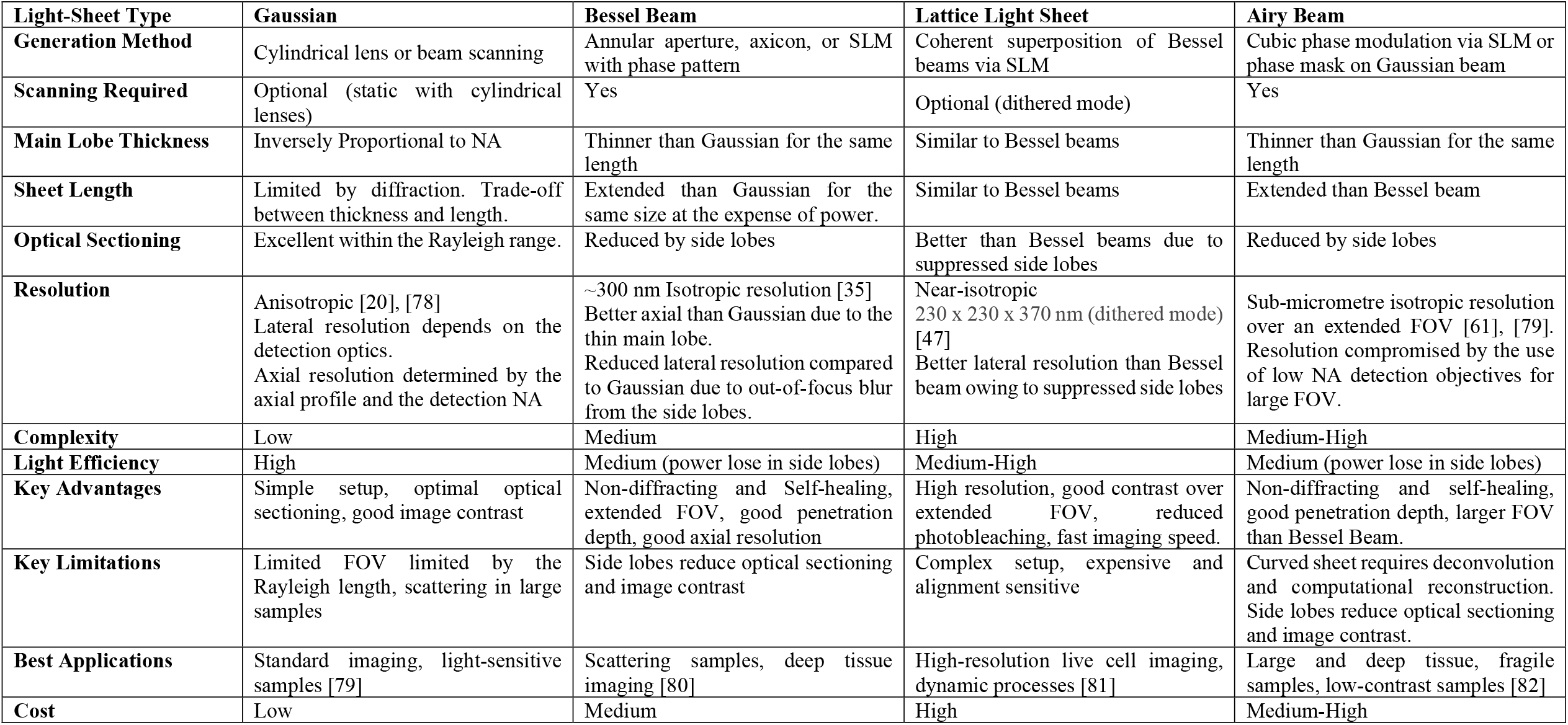
Comparison between different illumination strategies.

| **Light-Sheet Type** | **Gaussian** | **Bessel Beam** | **Lattice Light Sheet** | **Airy Beam** |
| --- | --- | --- | --- | --- |
| **Generation Method** | Cylindrical lens or beam scanning | Annular aperture, axicon, or SLM with phase pattern | Coherent superposition of Bessel beams via SLM | Cubic phase modulation via SLM or phase mask on Gaussian beam |
| **Scanning Required** | Optional (static with cylindrical lenses) | Yes | Optional (dithered mode) | Yes |
| **Main Lobe Thickness** | Inversely Proportional to NA | Thinner than Gaussian for the same length | Similar to Bessel beams | Thinner than Gaussian for the same length |
| **Sheet Length** | Limited by diffraction. Trade-off between thickness and length. | Extended than Gaussian for the same size at the expense of power. | Similar to Bessel beams | Extended than Bessel beam |
| **Optical Sectioning** | Excellent within the Rayleigh range. | Reduced by side lobes | Better than Bessel beams due to suppressed side lobes | Reduced by side lobes |
| **Resolution** | Anisotropic [20], [78]<br>Lateral resolution depends on the detection optics.<br>Axial resolution determined by the axial profile and the detection NA | ~300 nm Isotropic resolution [35]<br>Better axial than Gaussian due to the thin main lobe.<br>Reduced lateral resolution compared to Gaussian due to out-of-focus blur from the side lobes. | Near-isotropic<br>230 x 230 x 370 nm (dithered mode) [47]<br>Better lateral resolution than Bessel beam owing to suppressed side lobes | Sub-micrometre isotropic resolution over an extended FOV [61], [79].<br>Resolution compromised by the use of low NA detection objectives for large FOV. |
| **Complexity** | Low | Medium | High | Medium-High |
| **Light Efficiency** | High | Medium (power loss in side lobes) | Medium-High | Medium (power loss in side lobes) |
| **Key Advantages** | Simple setup, optimal optical sectioning, good image contrast | Non-diffracting and Self-healing, extended FOV, good penetration depth, good axial resolution | High resolution, good contrast over extended FOV, reduced photobleaching, fast imaging speed. | Non-diffracting and self-healing, good penetration depth, larger FOV than Bessel Beam. |
| **Key Limitations** | Limited FOV limited by the Rayleigh length, scattering in large samples | Side lobes reduce optical sectioning and image contrast | Complex setup, expensive and alignment sensitive | Curved sheet requires deconvolution and computational reconstruction. Side lobes reduce optical sectioning and image contrast. |
| **Best Applications** | Standard imaging, light-sensitive samples [79] | Scattering samples, deep tissue imaging [80] | High-resolution live cell imaging, dynamic processes [81] | Large and deep tissue, fragile samples, low-contrast samples [82] |
| **Cost** | Low | Medium | High | Medium-High |

### Light sheet scanning methods for high-speed volumetric imaging

Light-sheet microscopy has become widely adopted for volumetric imaging owing to its low phototoxicity and minimal photobleaching. Typically, volumetric imaging is achieved by translating the sample through the light sheet plane. To minimise mechanical perturbation of the sample and to improve imaging speed, remote refocusing was first introduced by Botcherby *et al*. [63]. The method is based on an optical configuration in which the overall lateral magnification, *M*_*RF*_, from the sample space (at the focal plane of objective O1) to the intermediate image space (at the focal plane of objective O2) is set equal to the ratio of the refractive indices of the two media by an additional pair of lenses. This condition ensures that the intermediate image is geometrically undistorted and free from refraction-induced aberrations. The intermediate image, generated by this pair of back-to-back 4f systems, is then re-imaged by a tertiary microscope for detection. This remote-refocusing configuration enables axial refocusing by translating the focal plane of the primary objective, without the need for active optical elements, i.e., components that dynamically alter optical properties such as refractive index or curvature to modulate light propagation [64], [65].

With the development of active optical elements, new approaches have emerged that utilise focus-adjustable devices such as electrically tunable lenses (ETLs) [66] and tunable acoustic gradient (TAG) lenses [67] for rapid refocusing. These methods achieve axial scanning by introducing low-order quadratic defocus, enabling fast volumetric imaging with low numerical aperture (NA) objectives. However, they are unable to maintain diffraction-limited performance in systems employing high NA objectives, where precise control of the wavefront is critical [68]. In such systems, refocusing via simple defocus introduces higher-order aberrations, particularly spherical aberration and field curvature, that degrade image quality and resolution across the imaging volume. To address this limitation, more advanced active optical elements, such as adaptive optics using deformable mirrors, have been integrated into light-sheet fluorescence microscopy (LSFM) for video-rate volumetric imaging [68]. Nevertheless, the implementation of these elements requires sophisticated hardware configurations and complex software-based wavefront calibration, substantially increasing system complexity and hindering broader adoption.

Although the remote-refocusing method proposed by Botcherby *et al*. does not actively correct for sample-induced aberrations (e.g., due to refractive index mismatches or inhomogeneities within the sample), its optical design, specifically satisfying the intermediate image magnification condition, allows for excellent control of spherical aberration during the refocusing process [63]. Consequently, this approach has seen widespread use in recent years.

With the continuous evolution of light-sheet microscopy, particularly following the invention of the single-objective light-sheet microscopy technique known as the OPM [12], a variety of scanning and refocusing strategies based on Botcherby *et al*.’s remote-refocusing architecture have emerged (see Table 2). These strategies can be broadly categorised into two types: scanning and reconstruction along the primary objective’s optical axis and scanning parallel to the primary objective’s focal plane, each offering distinct advantages.

**Table 2:**
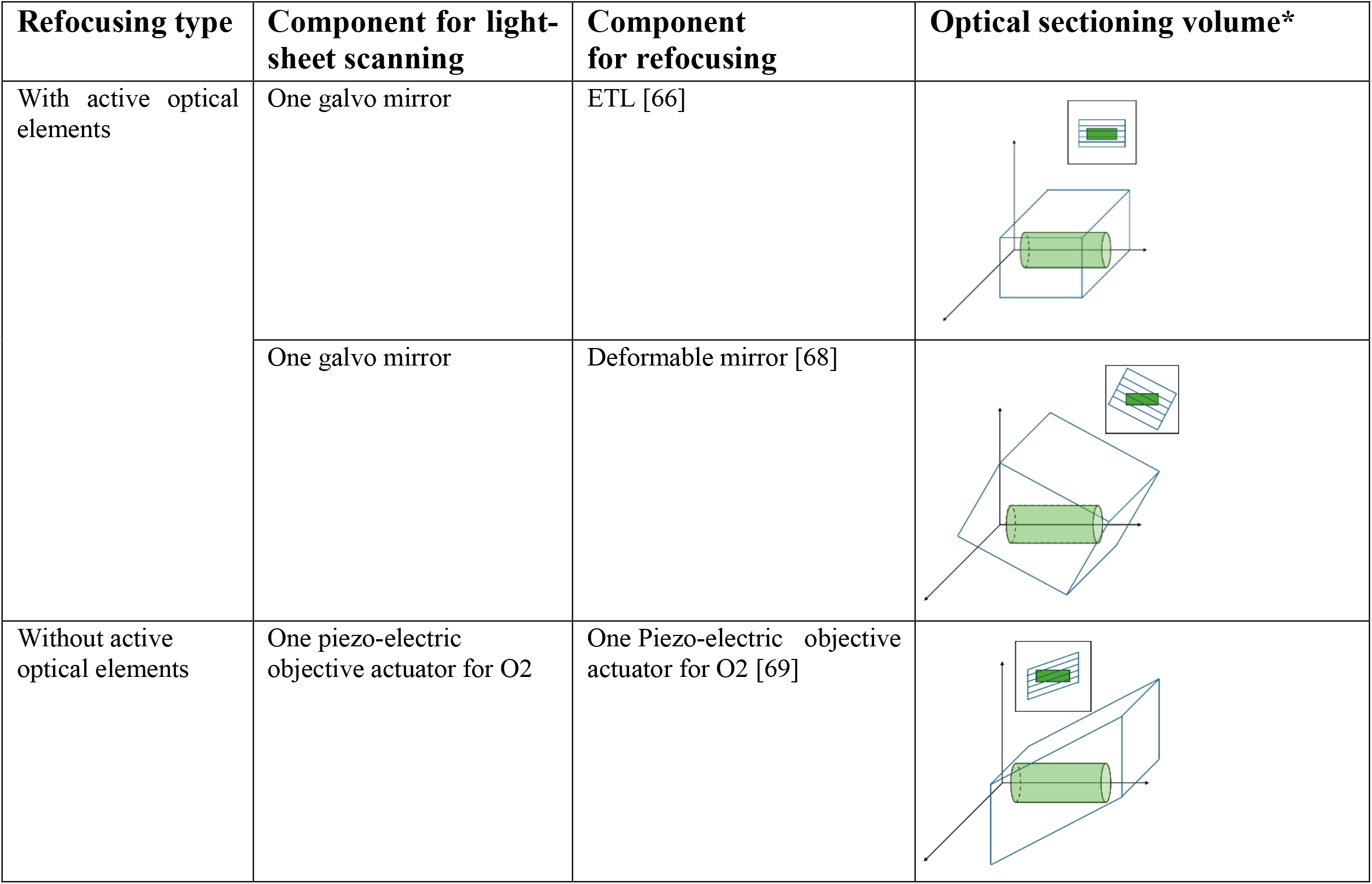
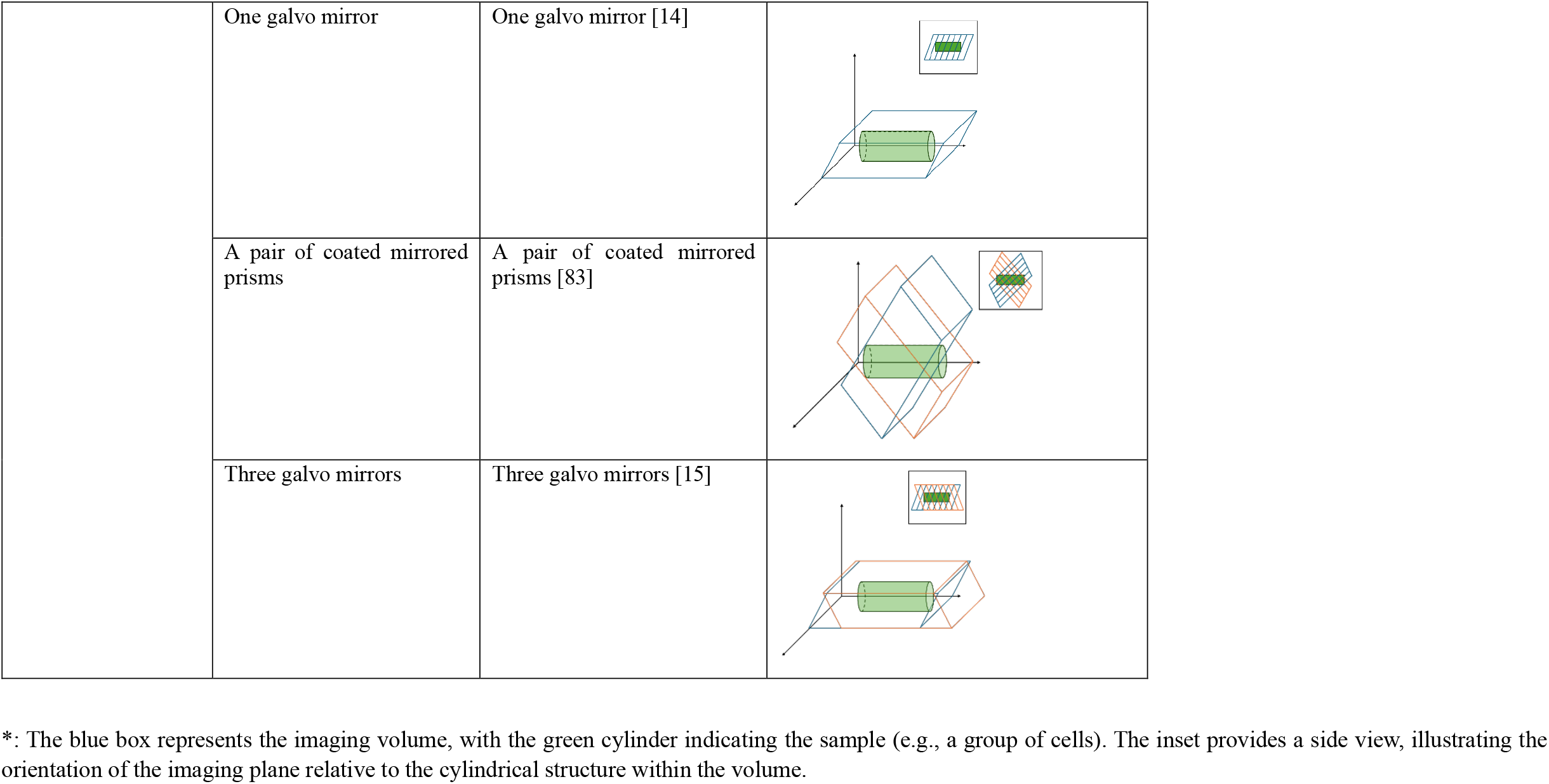
Methods of light-sheet scanning and refocusing for volumetric imaging.

| Refocusing type | Component for light-sheet scanning | Component for refocusing | Optical sectioning volume* |
| --- | --- | --- | --- |
| With active optical elements | One galvo mirror | ETL [66] |  |
|  | One galvo mirror | Deformable mirror [68] |  |
| Without active optical elements | One piezo-electric objective actuator for O2 | One Piezo-electric objective actuator for O2 [69] |  |
|  | One galvo mirror | One galvo mirror [14] |  |
|  | A pair of coated mirrored prisms | A pair of coated mirrored prisms [83] |  |
|  | Three galvo mirrors | Three galvo mirrors [15] |  |

\*: The blue box represents the imaging volume, with the green cylinder indicating the sample (e.g., a group of cells). The inset provides a side view, illustrating the orientation of the imaging plane relative to the cylindrical structure within the volume.

Axial scanning (along the optical axis) utilises the aberration-free refocusing capability of the remote-refocusing structure, but typically suffers from a limited lateral field of view, thereby requiring sample-stage scanning to cover larger areas [69]. In contrast, lateral scanning (parallel to the focal plane) enables the capture of a wider field of view in a single scan but limits axial imaging performance, often requiring axial repositioning of the objective to access deeper sample regions [14]. Recently, dual-view scanning strategies of each approach have also been developed to enhance resolution and volumetric detail [15], [70].

In this work, to achieve a larger lateral imaging area while maintaining high imaging speed, we focus on the construction and implementation of an OPM system employing lateral scanning parallel to the objective’s focal plane. We detail the specific design considerations and system setup involved in achieving efficient image acquisition with this configuration.

### System setup

Figure 2 shows the schematic and optical layout of the single-objective OPM system, with a full component list provided in Table 3. The system adopts a compact two-layer design: the bottom layer contains the excitation path, while the upper layer contains the emission path. The main setup only occupies approximately 85 cm × 70 cm × 45cm.

**Figure 2:**
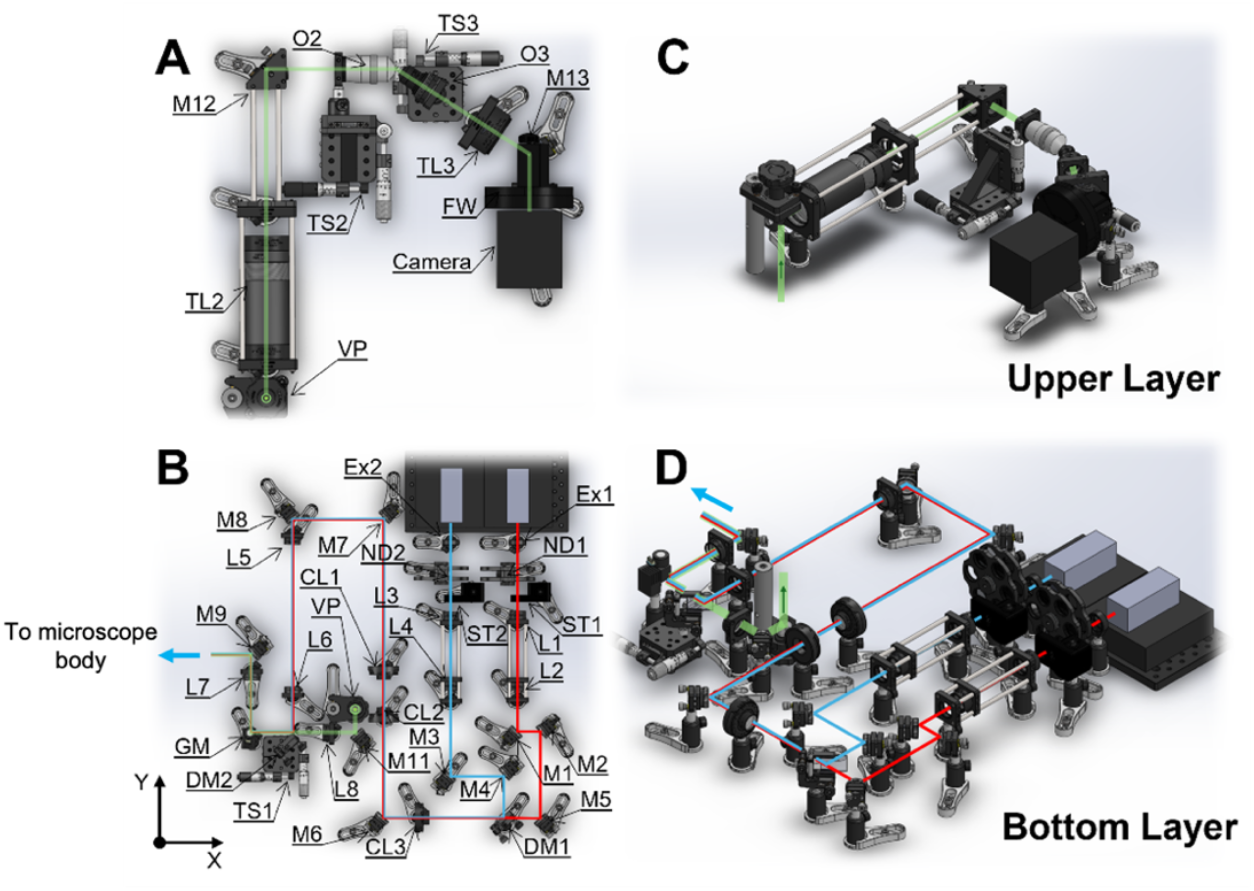
Optical layout of the OPM system used in this protocol. The system comprises two distinct optical layers. (A) Top-view schematic of the upper layer, showing the emission path. (B) Top-view schematic of the bottom layer, detailing the common path and excitation path. (C, D) 3D perspective views of the upper and bottom layers, respectively, to better illustrate the spatial arrangement of optical components. O, objective; L, lens; M, mirror; TL, tube lens; FW, filter wheel; DM, dichroic mirror; CL, cylindrical lens; VP, vertical periscope; TS, translation stage; Ex, excitation filter; GM, galvanometric mirror; ST, optical beam shutter; ND, neutral density filter; EF, emission filter.

The optical design centres around a 100× high-NA silicon oil immersion objective (O1) used for both light sheet excitation and fluorescence collection. The excitation path on the bottom layer includes two fixed-wavelength diode lasers (488 nm and 638 nm), with space for additional lines if needed. Each laser passes through its corresponding excitation filter and a logarithmic ND filter for intensity control. Shutters on each line enable independent beam control. Each laser beam is collimated by a beam expander (two plano-convex lenses). The laser lines are then combined via DM1 and focused by a cylindrical lens (CL1) into a 1D Gaussian sheet. This sheet can be expanded in width by inserting a cylindrical lens pair (CL2 & CL3) between DM1 and CL1. The beam is further reflected by a quad-band dichroic mirror (DM2), a 1D Galvo mirror (GM) and a mirror M9 into the microscope’s right camera port. The light-sheet tilt is controlled by translating DM2 along the optical axis.

Fluorescence collected by O1 is relayed to a 40× air objective (O2) on the upper layer of the system via two 4f systems (TL1 & L7 and L8 & TL2), aligned to conjugate the pupil planes of O1 and O2. It should be noted that for a commercial microscope such as the Nikon Ti series, the TL1 is incorporated into the main microscope body. The remote-refocusing module consists of a custom glass-tipped tertiary objective (O3) set at a 35° tilt, a tube lens (TL3), a motorised filter wheel (FW) for multi-colour imaging, and an sCMOS camera.

The system is built on a commercial inverted microscope body (Eclipse TE2000-U, Nikon). O1 is mounted on the nosepiece, beneath a 2D motorised stage with custom sample holders. The GM between L7 and L8 adjusts the lateral position of the light-sheet, positioned to be conjugate to both O1 and O2 pupil planes. The rotation angle of the GM was modulated by the input voltage generated by a digital-to-analogue converter (DAC), which was controlled by a home-written LabView program. This enables light sheet scanning at a speed limited by the camera frame rate. In this design, the GM was the only moving element in the entire optical setup, which minimised any mechanical drift and thereby augmented the stability of the OPM system.

### Designing Your Own OPM Microscope: theoretical considerations

This guide outlines the key design considerations for building an OPM system employing a Gaussian light sheet and an RFS, highlighting the selection of optical components for cellular imaging. It should be emphasised that the specific design of the OPM system, including the choice of the objective lens(es) and other optical components in the emission path, primarily depends on the selected primary objective, which should be carefully chosen to suit designated imaging tasks. An excel file “OPM_Easy2Use” is provided to facilitate researchers wishing to design their own OPM system.

### Step 1: Determine the Light-Sheet Characteristics

#### 1.1 Calculate the Depth of Field (DOF)

Optimal optical sectioning in LSFM suggests the beam waist of the light sheet to match the DOF of the primary objective O1 [71]:

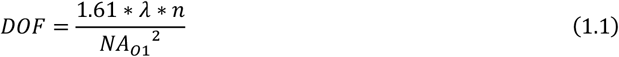

λ: illumination wavelength.

*n*: refractive index of the immersion medium.

*NA*_*o1*_: numerical aperture of the primary objective O1.

- **Example:** For a 100×, 1.35 NA silicone oil objective at *λ*= 638 nm and *n* = 1.4, DOF ≈ 0.79 µm.

#### 1.2 Determine the Rayleigh Length

Rayleigh length (*z*_*R*_) affects the imaging volume and is calculated by:

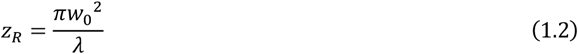

*w*_*o*_: beam waist of the Gaussian light sheet.

- **Example:** A beam waist of 0.79 µm yields a Rayleigh length of 3.07 µm with *λ*= 638 nm.
- **Consideration:** While thinner sheets improve axial resolution, they shorten the Rayleigh length, thus reducing the imaging depth.

#### 1.3 Calculate Imaging Depth (*D*_*im*_)

Considering the light sheet tilt *θ*_*tilt*_:

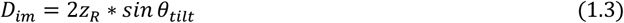

- **Example:** For a 35° tilt of the light sheet and an imaging depth of ~10 µm, the beam waist *w*_0_ ≈ 1.33 µm.
- **Consideration:** While thinner sheets improve axial resolution, they shorten the Rayleigh length and reduce the imaging depth. The tertiary objective (O3) can tilt between 0° and 45°. Increasing the tilt angle helps extend the imaging depth but reduces the overall detection NA. (Figure S1: NA calculation)

#### 1.4 Choose the Final Beam Waist

Select a beam waist that balances the resolution and imaging depth. Here, 1.33 µm was chosen to ensure an axial resolution sufficient for subcellular imaging.

### Step 2: Select Primary Objective (O1)

For cellular imaging, a high NA objective is ideal due to its ability to produce a thin light-sheet, collect light efficiently and achieve higher spatial resolution.

- **Example:** 100×, 1.35 NA silicone oil objective (RI ≈ 1.4) reduces the refractive index mismatch with aqueous samples (RI ≈ 1.38), reducing spherical aberrations common with water immersion objectives [72].
- **Alternative options:** Lower magnification objectives (e.g., Olympus 20× XLUMPLFLN, 1.0 NA) can be chosen for a larger FOV.

### Step 3: Select Secondary Objective (O2)

O2 collects the fluorescence emission and maps the pupil plane of O1. The angular apertures of O1 and O2 should be matched for light collection efficiency. The angular aperture *θ* can be calculated from:

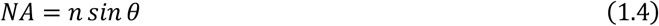

- **Example:** For a 100×, 1.35 NA silicone oil objective, the angular aperture = 74.6°
- **Chosen:** 40×, 0.95 NA air objective with a similar aperture of 71.8°.

### Step 4: Remote-refocusing Range Calculation

In the remote-refocusing system, once the parameters of Microscope 1 (O1+TL1) and Microscope 2 (O2+TL2) are set, the system’s refocusing capability becomes fixed. This is quantified as the geometric refocus distance *Z*_*max*_, as defined in [73]:

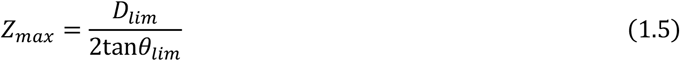

*D*_*lim*_ is given by the minimum of the field number of O1, *FN*_*o1*_, and the field number of O2, *FN*_*o*2_, projected into sample space, i.e., 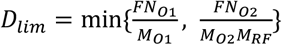. Meanwhile, *θ*_*lim*_ is given by the minimum angular acceptance of O1 and O2, i.e., 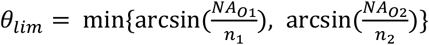

- **Example:** For a remote-refocusing system with a 100×, 1.35 NA silicone oil objective (O1, *FN*_*o1*_ = 25 mm), and a 40×, 0.95 NA air objective (O2, *FN*_*o*2_ = 25 mm), *θ*_*lim*_ = 71.8°, *D*_*lim*_ = 250 μm, *Z*_*max*_ = ~41μm. This means that this system is able to achieve refocusing of ~82 µm covering both sides of the focal plane of O1.

### Step 5: Select Tertiary Objective (O3)

- **Chosen:** AMS-AGY v1.0 (1.0 NA glass-tipped) for improved emission collection.
- **Advantages:**
  - High fluorescence collection efficiency due to high NA.
  - The glass tip introduces a refractive index mismatch at the air-glass interface which compresses the emission light cone for better collection efficiency.
  - Zero working distance eliminates emission loss.
  - Anti-reflection coatings mitigate spherical aberrations.
  - **Alternative option:** AMS-AGY v2.0 for larger FOV imaging.

### Step 6: Select Primary Tube Lens (TL1)

TL1 is often integrated into the microscope body and cannot be changed.

- **TL1:** 200 mm infinity-corrected tube lens in the Nikon microscope body.
- TL1 may have different focal lengths across manufacturers. For example, the built-in tube lens in Olympus microscope bodies has a focal length of 180 mm, while in Zeiss microscopes it is 165 mm.

### Step 7: Select Scan Lenses (L6 & L7 & L8)

In OPM, volumetric imaging is achieved by rapidly scanning the light-sheet through the specimen with a 1-D motorised galvo mirror. Unity magnification at the Galvo mirror is required, suggesting:

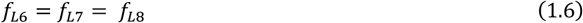

The scan lenses should accommodate the range of laser wavelengths and the size of the beam, *d*_*SL*_:

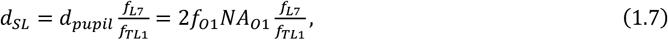

Where

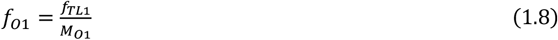

*d_pupil_*: pupil diameter of O1.

*f*_*o*1_: focal length of O1.

*M*_*o*1_: magnification of O1.

- **Chosen:** 100 mm achromatic lenses.
- **Example:** Beam diameter (*d*_*SL*_) ≈ 2.7 mm, accommodating standard 1” optics.

### Step 8: Select Galvo Mirror

To avoid clipping, the size of the beam at the galvo mirror, *d*_*galvo*_, should satisfy:

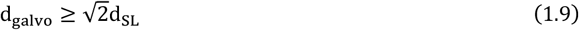

- **Example:** *d*_*galvo*_ ≈ 3.82 mm.
- **Chosen:** 5 mm diameter 1D galvo scanner (GVS201, Thorlabs).

### Step 9: Select Secondary Tube Lens (TL2)

The magnification of the intermediate image at the focal plane of O2, *M*_*RF*_, must match the refractive index of the immersion medium of O1 to minimise aberration. This magnification directly influences the choice of lenses in the two 4f systems (TL1-L7 and L8-TL2). Given selected O1 and O2, the required effective focal length (EFL) for TL2 is:

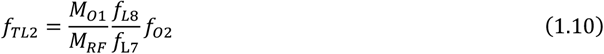

- **Implementation:** A low-cost tube lens assembly consists of commercially available achromatic lenses and lens tubes [74]. The separation distance *d* between the lenses for the tube lens assembly that yields the desired EFL (*f*_*EFL*_) can be determined by:

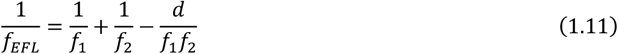

*f*_1_ and *f*_2_: focal lengths of the lenses used for the tube lens assembly.
- **Example:** For a 100×, 1.35 NA silicone oil O1 and a 40×, 0.95 NA air objective O2 (*f*_*o*2_ = 5 mm), *f*_*TL*2_ ≈ 357 mm.
- This can be achieved with a lens pair (500 mm & 1000 mm achromatic doublets) with a 100.3 mm edge separation (Figure S4). Standard 2” optics is required to accommodate the emission beam size and the scan range.

### Step 10: Expand Beam for Light Sheet Generation (L1 & L2, L3 & L4)

To control the light-sheet thickness and length, adjusting the beam size entering the primary objective (O1) is essential. High NA objectives can produce thinner light sheets but at the cost of shorter propagation distances. To achieve a slightly thicker yet longer light sheet, the beam size at the back focal plane (BFP) of the objective must be smaller than the full aperture of O1. This reduces the effective NA used for illumination, allowing a balance between sheet thickness and imaging depth.

The effective light-sheet NA can be approximated as (for small *θ*):

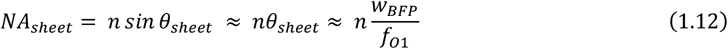

*w*_*BFP*_: general beam waist of the input gaussian beam at the BFP of the objective.

For light-sheet with a beam waist *w*_0_, the radius at the BFP of the objective along the non-focused axis, *r*_*obj*_, is equivalent to *w*_*BFP*_:

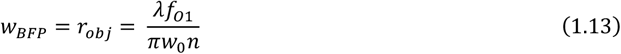

- **Example:** For *w*_0_ = 1.33 μm at 638 nm and n = 1.4, *r*_*obj*_ ≈ 218.1 um.

Commercial laser sources typically provide a beam radius of ~350 µm (at 1/e^2^). To manage this size and correct the inherent beam divergence, a 1× beam expander (collimator) was selected, ensuring the beam size remains suitable for the desired light-sheet characteristics:

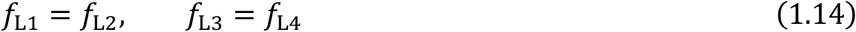

This symmetrical configuration preserves the beam size while improving collimation. An adjustable iris can be placed at the focal plane of CL1 to further refine the beam diameter.

### Step 11: Determine the Light Sheet Width

#### 11.1 Select Focusing Lens (L5)

Given the size of the beam at the laser source, *r*_*laser*_, the radius of the beam at the BFP of O1, *r*_*obj*_, is also related to the focal length of L5:

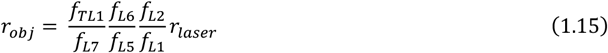

- **Example:** For *r*_*obj*_ = 218.1 μm and *r*_*laser*_ = 350 μm, L5 ≈ 320.9 mm.
- **Chosen lens:** 300 mm achromatic lens.
- The longer focal length compensates for divergence and ensures a stable beam waist at the back focal plane of O1, producing a well-defined light sheet.

#### 11.2 Select Cylindrical Lens (CL1)

The choice of CL1 determines the width of the light sheet, and therefore the imaging FOV. The width of the light sheet can be calculated as [75]:

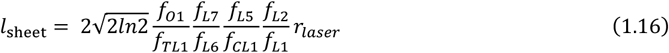

- **Example:** A 100 mm CL1 produces a 24.7 µm-wide light sheet suitable for standard cellular imaging.
- **Alternative option:** A 50 mm cylindrical lens for a broader FOV.

### Step 12: Select Camera

The camera chip must be sufficiently large to accommodate the FOV of O3. The number of pixels (*n*_*pixel*_) required to cover this FOV is related to the Nyquist resolution *R*_*nyquist*_ of the system and can be determined by [76], [77]:

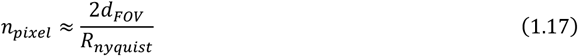

where the Nyquist resolution is determined by:

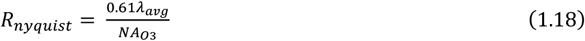

*d*_*FOV*_: diameter of the diffraction limited FOV.

*NA*_*o*3_: NA of the AMS-AGY v1.0 objective.

The theoretical lateral resolution limit of the system, can be estimated from the system NA and wavelength:

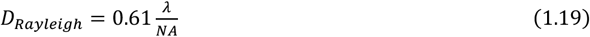

- **Example:** For *λ*_*avg*_ = 550 nm and *NA*_*o*3_ = 1, *R*_*nyquist*_ ≈ 335.5 nm.
- AMS-AGY v1.0 provides a 150 μm diameter of a diffraction-limited FOV of 150 µm, *n*_*pixel*_ ≈ 894 pixels.
- A high quantum efficiency (QE), low read noise sCMOS camera is recommended. For fast live volumetric imaging, a high frame rate is also desirable.

### Step 13: Select Tertiary Tube Lens (TL3)

The focal length of the tertiary tube lens, *f*_*TL*3_, is related to the Nyquist resolution:

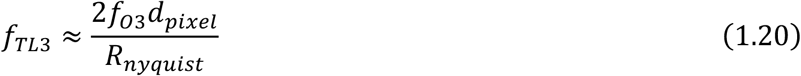

*f*_*o*3_: focal length of O3.

*d*_*pixel*_: physical pixel size of the camera.

- **Example:** To satisfy the Nyquist resolution with *f*_*o*3_ = 5 mm and *d*_*pixel*_ = 6.5 um, *f*_*TL*3_ = 193.7 mm.
- **Chosen lens:** 200 mm infinity-corrected tube lens.

### Step 14 (Optional): Extend the light sheet width

A pair of cylindrical lenses can be placed before CL1 to further expand the beam size along the width of the diaphragm, but at the cost of lost power density. Depending on the desired width, 50 mm (CL3) × 100/150/200 mm (CL2) is a good choice.

### Alignment procedure

This section outlines the alignment procedure for a single laser line (indicated as the 488 nm blue laser line in Figure 2) with a commercial inverted microscope body (Eclipse TE2000-U, Nikon). After successfully aligning the first laser line, the alignment process for additional laser lines can follow the same procedure.

A detailed part list and a complementary assembly and alignment guide containing annotated photos to aid the system construction is also included as the supplementary material.

### General requirements

Required space on the optical table including the microscope body and the reference path: 110 × 85 x 40 cm. Optical breadboard (Metric): mounting hole spacing 2.5 cm.

A dichroic cube inside the microscope body is not required.

The 1.5× magnification lens built-in the microscope body should not be used.

### Pre-requisite: Two-iris Alignment Technique

Using two irises to define the optical axis is a fundamental alignment technique that is extensively used in this protocol. An iris, or iris diaphragm, is a circular optical component with an adjustable aperture located at its centre. By using two coaxial irises with the same height in the optical path, one can ensure that the laser beam is level with respect to the surface of the optical table. This method ensures precise alignment of optical components and maintains a well-defined beam path throughout the system.

#### 1. Placement of Irises

- Position two irises (ID25/M) with post holders (PH40/M, Thorlabs) along the intended optical path with a sufficiently large separation (e.g., ~200 mm) between them.
- Where possible, mount the post holders directly onto the optical table by securing them into pre-drilled mounting holes without using bases or clamping forks. This ensures that the iris centre is right above the mounting hole.
- Mount the two irises onto the same row of holes on the optical table to ensure that the optical axis is strictly parallel to the row of holes.

#### 2. Beam Alignment

- Adjust the optical component (e.g., mirror, lens, dichroic) so that the beam passes through the centres of both irises simultaneously.
- If the beam does not pass through both centres, fine-tune the height, orientation, and/or position of the optical component until alignment is achieved.

#### 3. Key Considerations

- The beam path should remain aligned with the intended optical axis, which is often level and parallel to the rows of mounting holes on the optical table. Introducing lenses or mirrors should not significantly alter the beam height or position.
- Some components, like Galvo mirrors, may introduce slight deviations in height or angle. These can be corrected by adjusting downstream optics rather than forcing the beam to align at an unusual angle.
- Once aligned, the beam path established by the two irises serves as a reference for all downstream optical elements.

This technique is systematically applied throughout the protocol. Whenever an instruction refers to “aligning using two irises”, follow this method to ensure optimal optical alignment.

### Reference Path (Time: 3h)

A collimated reference laser beam is critical throughout the alignment process, as it defines the optical axis and helps position optics in both the illumination and detection paths.

1. (Photo 1A) in the alignment guide) Place the microscope body on a stable and level optical table. Reserve ~15 cm to the left for mounting the reference laser and ~80 cm to the right for constructing the main system.
2. (Photo 1B-E) Mount the collimated reference laser (CPS635R, Thorlabs) with a kinematic collimator adaptor (KAD11F, Thorlabs) approximately 15 cm above the sample stage with a 50 cm tall and 1.5’’ diameter post (2 × P250/M, Thorlabs) **Note:** The sample stage is assumed to have been correctly installed for this protocol.
3. (Photo 2A-E) Mount a threaded iris (SM1D12D, Thorlabs) onto one nosepiece port using an SM1-to-C-mount adapter (SM1A10, Thorlabs).
4. (Photo 2F-G) Mount another threaded iris of the same model with a 4.00’’ lens tube (SM1L40, Thorlabs) onto the adjacent nosepiece port using the same adapter. The centres of these two irises define a straight vertical optical axis for all downstream alignments.
5. (Photo 3A-D) Switch on the reference laser and adjust its position and tilt with the collimator adaptor so that the beam passes through the centres of both irises simultaneously.
6. Swap between the two nosepieces frequently to verify alignment. **Critical**: Passing the laser through the centre of both irises is critical, as it defines the optical axis for all subsequent optics in both excitation and emission paths. **Caution**: Ensure there is enough space when swapping nosepieces to prevent the extended lens tube from hitting the stage.
7. (Photo 3E) Mount the primary objective O1 onto a spare nosepiece port. The reference laser should also pass through the centre of the objective aperture. **Note:** The reference laser provides an approximation of the collimated emission line but does not perfectly represent the true emission. Some slight deviation when adding or removing O1 is expected.
8. Make sure the magnification of the microscope body is set to 1×.
9. (Photo 3F) Switch to the R port on the microscope body. A focused laser beam should be visible from the right camera port of the microscope body without O1.
10. With the reference path aligned, proceed with aligning the common path of the main light sheet system. **Safety Precaution**: The reference laser may be near eye level, so always wear appropriate laser safety goggles (e.g., LG4, Thorlabs) during alignment.

### Common path (3 hours)

The common path is shared by both the illumination and emission paths. It consists of a 100 mm achromatic lens L7 (AC254-100-A-ML, Thorlabs), a 1D Galvo mirror for rapid light sheet scanning, and a quad-band dichroic mirror (Di01-R406/488/561/635-25×36, Semrock) that separates the excitation and emission paths. The dichroic mirror is mounted on a 1D translation stage TS1 (LX10/M, Thorlabs), allowing precise tilting of the light sheet by translating the stage. The alignment procedure follows a backward approach, starting from the microscope body and working toward the laser source along the illumination path.

11. Identify the height for the excitation optics (in the bottom layer) by determining the optical axis at the right camera port. The optical axis should be level if the reference laser is vertical and passes through the centre of O1.
12. (Photo 4A) Place a post-mounted iris (without the base) as close to the camera port as possible. Adjust the height of the aperture so that the reference beam passes through its centre.
13. Repeat step 12 for another iris, and additional irises, if necessary, to confirm the height of the optical axis.
14. (Photo 4B) Position a mirror (M9) approximately at the focal point of the reference laser so that it reflects the focused beam by 90 degrees.
15. (Photo 4B) Use the Two-iris Alignment Technique to ensure the reflected beam follows a well-defined optical axis. The reflected beam should be diverging.
16. (Photo 4C) Mount a threaded iris in front of an achromatic lens L7 and place it along the optical path so that the beam passes through the centre of the lens and emerges collimated after L7.
17. (Photo 4D) Verify collimation with a shear interferometer (SI035, Thorlabs, 1–3 mm shear plate). Use a magnifier (SIVS, Thorlabs) for better visualisation of the fringes. **Critical**: When introducing any achromatic lens into the optical path, make sure that its placement should not alter the beam path through the irises. If necessary, adjust lens positioning to maintain alignment.
18. Switch on the Galvo mirror and set the voltage to 1.5 V to fix it at its default angular position. (Refer to Supplementary Information (SI) for details on connecting the Galvo mirror). **Critical:** Make sure the Galvo mirror is always turned on and fixed at the same angular position throughout the alignment.
19. Rotate the objective nosepiece and place O1 in the optical path. After passing through L7, the collimated beam should focus at a point roughly 100 mm downstream.
20. Before proceeding, ensure that objective O1 is positioned at the **correct imaging height** as follows. Set the correction collar of O1 to 0.17, which corresponds to the thickness of a #1.5 glass coverslip.
21. Use a black marker to draw a few small dots on the top surface of a #1.5 glass coverslip and place the coverslip on the sample stage.
22. Turn off the reference laser. Apply a drop of silicone immersion oil (MXA22179, Nikon) to the top of O1.
23. Set the optical path selector to “EYE” instead of “R”. **Caution:** Make sure the alignment laser is OFF before staring into the eye piece.
24. Adjust the height of O1 using the manual focus knobs on both sides of the microscope body until the black dots are clearly in focus when viewed through the eyepiece.
25. Once the dot is sharply focused, fix O1 at this height as it will serve as the reference imaging plane for all subsequent alignments. **Critical:** Determining the correct imaging height of O1 is crucial, as this height corresponds to the focal plane at the coverslip. This step also ensures that the optical distance between O1 and TL1 is correct, which is necessary for alignment of the magnification of the downstream 4f optical systems.
26. Discard the coverslip, remove the immersion oil from O1, turn on the reference laser, and switch the optical path selector back to ‘R’ for further alignment steps.
27. (Photo 4E-F) Position the Galvo mirror (GM) at the beam focus. A small, focused spot should appear at the centre of the mirror, and the beam should bend 90 degrees to the right and defocus as it propagates. **Critical**: Ensure the focus spot is as small as possible on the Galvo mirror. Fine-tuning may be needed later when setting up the emission line.
28. Use the Two-iris Alignment Technique to align the Galvo mirror so that the beam reflected by the GM passes through the centres of both irises. **Note:** The optical axis may not align directly with a row of holes on the optical table. Place the alignment irises accordingly so that the optical axis remains perpendicular to its previous direction. **Caution**: Since the 1D Galvo mirror has no tilt control, slight beam height variations may occur at longer distances. Downstream optics aligned with beam passing through the components’ centre will compensate for this, so perfect horizontal alignment is not required.
29. (Photo 5A) Secure the quad-band dichroic mirror in a rectangular optics holder (KM100C, Thorlabs) and mount it onto a 1D translation stage TS1.
30. (Photo 5B) Orient TS1 such that it moves along the X-axis of the optical table.
31. (Photo 5B) Position the dichroic mirror (DM2) roughly 5 cm from Galvo mirror to reserve some space for mounting lens L8.
32. (Photo 5C) The dichroic mirror should reflect most power of the beam 90 degrees along the Y-axis. A dim transmitted beam should continue along the X-axis, which will be used for aligning the emission path.
33. Use the Two-iris Alignment Technique again to align the beam along this reflected new optical path along the Y-axis.
34. Swap O1 with an adjacent empty nosepiece port. The collimated beam should now be reflected by both the Galvo and dichroic mirrors.
35. Adjust TS1 and the pitch knobs on the dichroic mirror mount to fine-tune the dichroic mirror position so that the reflected beam is centred at both irises along the Y-axis as shown in Photo 5C. **Caution**: Ensure the 1D translation stage TS1 is centred within its range, not at the extremes.
36. The common path alignment is now complete.

### Excitation Path (Part I: 1 hour)

37. (Photo 6A) Rotate the objective nosepiece and place O1 back into the optical path. Position a 100 mm achromatic lens L6 (AC254-100-A-ML, Thorlabs) at a position such that the beam becomes collimated after this lens.
38. Verify collimation using a shear interferometer (1-3 mm beam shear plate), ensuring that the lens does not alter the beam’s propagation direction.
39. (Photo 6B) Swap O1 with a spare nosepiece port again. A collimated beam should still be visible before entering L6. Introduce another achromatic lens L5 (AC254-300-A-ML, Thorlabs) approximately 40 cm further down the optical path so that the beam exiting L5 remains collimated. Check beam collimation with a shear interferometer (2.5-5 mm beam shear plate) and confirm that the lens positioning does not alter the beam’s orientation.
40. (Photo 6C) Place a mirror M8 just after L5 to reflect the beam 90 degrees to the right along the X-axis. Using the Two-iris Alignment Technique to align the beam along the new optical axis.

### Emission Path (Part I: 3 hours)

Before aligning the rest of the excitation path, the approximate position of the upper layer must be determined. This ensures that the optics in the lower excitation path can be positioned correctly without being obstructed by the posts supporting the upper layer.

41. (Photo 7A) Fixed three 200 mm posts (P200/M) to support the upper layer. The right top corner is not supported by a post, as the lasers will be mounted under that area.
42. (Photo 7A) Position the upper layer so that the left edge of the breadboard aligns with the right edge of the 1D translation stage TS1. Maintain a gap of approximately 4-5 holes (~10 cm) from the top edge of the 1D translation stage TS1. Do not secure the posts yet, as fine adjustments will be required during emission path alignment.
43. (Photo 7B) Bring O1 back into the optical path. The reference beam should be focused at the Galvo mirror. Use the weak transmitted reference beam through the dichroic mirror to align the downstream emission path. Switch off the room lights if needed to better visualise the weak transmission.
44. (Photo 7C-D) Similar to the alignment of L6, use the weak transmitted beam to align L8 (AC254-100-A-ML, Thorlabs), ensuring that the beam exiting the lens is collimated with the shear interferometer (1-3 mm beam shear plate).
45. (Photo 7E) Position a mirror (M11) approximately 5 cm after L8 to reflect the beam 90 degrees.
46. (Photo 7E) Assemble the vertical periscope (VP) system (RS99/M, Thorlabs) and place it approximately 5 cm down the Y-axis after M11.
47. Ensure that the upper-layer breadboard aligns with the mounting hole grids of the lower optical table. The breadboard should not be rotated at an angle relative to the lower table, and the posts supporting the upper breadboard must not obstruct L6 in the excitation path.
48. Place two irises at a distance apart along the third column of hole on the breadboard to define the optical axis of the upper layer. The centre of each iris should be approximately 10 cm above the breadboard. **Note:** This step defines the optical axis for the upper layer.
49. Adjust the three mirrors (M11 and VP) so that the collimated beam passes through the centres of both irises.
50. Adjust both the orientation of the bottom mirror and the height of the upper mirror in the vertical periscope for fine alignment.
51. Tune the angle of M11 to centre the beam on the bottom VP mirror and tune the orientation of the bottom VP mirror to centre the beam on the upper VP mirror.
52. Tune the upper VP mirror’s rotation angle and its tip-tilt to centre the beam through the irises. These alignment steps may require a few iterations. **Critical**: The beam must hit the centres of both the bottom and upper mirrors of the vertical periscope. Off-centre misalignment can result in clipped FOV in the final images.
53. Once the positions of M11 and the vertical periscope are confirmed, the position of the upper layer is finalised. Secure the posts of the upper layer onto the optical table with clamps.

### Excitation Path (II: 4 hours)

54. (Photo 8A) Swap O1 with a spare nosepiece port again and place a mirror (M7) approximately seven holes (17.5 cm) away from M8 to reflect the beam 90 degrees along the Y-axis. Ensure that the reflected beam does not clip on M11 or the vertical periscope.
55. Align the excitation path forward from the laser head. The goal is to overlap the laser beam with the reference beam to ensure optimal alignment.
56. (Photo 8B) Mount the lasers with their heatsinks onto a 12.7 cm thick breadboard (MB1530/M) to ensure that the height of the beam approaches the pre-determined height of the optical axis.
57. (Photo 8B) Mount the entire setup onto the optical table beneath the upper layer. Ensure that the 638-nm laser beam is not blocked by the posts supporting the upper layer. There should be sufficient space for mounting the beam expander, shutter, and mirrors, so the exact positioning of the laser heads is not critical.
58. Connect and switch on the 488-nm laser at low power (2-5 mW) for safety.
59. (Photo 8B) Place the 488-excitation filter (Ex2, MF475-35, Thorlabs) and the neutral density filter wheel (FW2AND, Thorlabs) sequentially along the excitation path.
60. (Photo 8C) Set up the beam expander (L3 and L4) in a cage system. Two SM1 cage plates (CP33/M, Thorlabs) should be connected using four 6” cage assembly rods (ER4-P4, Thorlabs) and two posts at both ends of the cage system.
61. The front cage plate (facing the laser head) will hold L3 (AC254-050-A-ML, Thorlabs).
62. A slidable SM1 cage in the middle will hold L4 (AC254-050-A-ML, Thorlabs), with an additional cage iris (CP20D, Thorlabs) to indicate the optical centre.
63. Mount a threaded iris (SM1D12D, Thorlabs) onto the front cage plate and introduce the cage system (without L3 and L4) into the optical path after the ND filter.
64. Adjust the position and height of the entire cage system so that the laser beam passes through the centre of both irises. **Note**: You can recycle the threaded irises from the nosepieces.
65. (Photo 8D-E) Remove the front iris, insert L3 and L4 into the cage system, then reposition the front iris in front of L3.
66. Adjust L4 along the cage system to produce a collimated beam and fix its position by tightening the screws on the sides of the cage plate. **Critical**: Since the expanded beam diameter is very small, a shear interferometer cannot reliably check collimation. Instead, check the beam collimation at a distance (>1 m) from L4 and adjust L4 until the beam spot appears the smallest.
67. Ensure that all beam expander components are at the same height and that the beam’s orientation remains **unchanged** with and without the lenses. **Critical**: The step assumes that the laser beam is parallel to the optical table, meaning the beam height should not change significantly.
68. (Optional) Mount a 100 μm pinhole (P100K, Thorlabs) onto an XY Translating Mount (CXY1A, Thorlabs) and insert it at the focal point of L3 to clean the beam and to achieve the TEM00 mode (the fundamental Gaussian beam with a smooth, single-lobed profile).
69. (Photo 9A) Position mirrors M3 and M4 sequentially after the beam expander. These mirrors form a periscope system to enable fine adjustments of the beam direction and correct any off-axis beam.
70. (Photo 9A) Mount a single-band dichroic mirror DM1 (Di02-R488-25×36, Semrock) on a kinematic dichroic mount (KM100C, Thorlabs) and place it after M4 to reflect the beam 90 degrees left along the X-axis.
71. Use the Two-iris Alignment Technique to check the propagation direction of the beam. Adjust M3 and M4 knobs to align the beam through both irises. **Critical**: Position the dichroic mirror so that the reflected beam does not interfere with emission optics (the Galvo mirror, TS1, L8 and M11), leaving sufficient space for two irises.
72. At this point, the reference beam and the excitation beam from the laser head should **intersect**.
73. (Photo 9A-B) Place a mirror M6 at the intersection point. Adjust the knobs until the two beams overlap completely. If necessary, fine-tune M3, M4, and M6 to ensure **full overlap**. **Critical**: (Photo 9C-D) It is essential that the two beams fully overlap at every point along the optical path. Insert an alignment checking card (e.g., a small piece of white paper) into the beam path and check for overlap, especially at the microscope nosepiece (O1 position).
74. (Photo 9E) Mount the optical beam shutter ST2 (SH05R/M, Thorlabs) after the beam expander (L3 & L4).
75. Bring O1 back into the path. Position a 100 mm achromatic spherical lens L9 (AC254-100-A-ML, Thorlabs) between M6 and M7 so that the reference beam is collimated before entering L5 and remains collimated after exiting L9. Use a shear interferometer (1-3 mm beam shear plate) to verify collimation. This L9 will later be replaced by a cylindrical lens CL1(ACY254-100-A, Thorlabs) for generating the light sheet. **Note**: An achromatic lens is used at this step for easier collimation checks with a shear interferometer.
76. With O1 in place, using the shear interferometer to verify that O1-TL1, L7-L6, and L5-L9 (CL1) each form a 4f system.
77. Temporarily turn off the reference beam. Move a small piece of paper towards and away from O1 to check the collimation, a ‘collimated’ beam exiting O1 should be observed. **Critical**: The exiting beam may not be perfectly collimated and might have a slight divergence. If the beam appears defocused as it moves away,
78. At this point, the beam should remain aligned with the optical axis if you move a small piece of paper along the optical axis. **Critical**: A slight off-axis shift of the exiting beam is normal. If the beam does not fully overlap with the reference beam at a distance, this is due to minor misalignment at the back focal plane of O1. This can be compensated when the cylindrical lens (CL1) is introduced in the next step.
79. (Photo 10A-B) Mount the cylindrical lens (CL1) onto a rotational mount (RSP1D/M) and use a threaded iris to help identify the centre of the cylindrical lens.
80. (Photo 10C) Switch the reference laser back on and replace L9 with the cylindrical lens (CL1). The curved surface should face towards the incoming excitation beam from the laser head.
81. (Photo 10D) Rotate CL1 along its centre until a sharp line along the Y-axis is seen at O1.
82. If the centre of the light sheet does not overlap with the reference laser, adjust CL1’s lateral position (perpendicular to the optical axis) and height until they align.
83. Verify that the two beams remain overlapped at many other positions in the illumination path.
84. Position an iris at the focal plane of CL1 to control the beam size and the sheet property.
85. Repeat Step 53-72 for aligning other excitation laser lines. All laser lines should overlap with the reference beam and each other.
86. The excitation path alignment is now complete.

### Emission path (II: 6 hours)

87. Continue the alignment from the VP system into the upper layer.
88. Keep O1 in the optical path, a collimated beam should pass through two irises of the same height (~10 cm above the table) along a straight line.
89. (Photo 11A) Assemble the secondary tube lens TL2 using the adjustable lens tube (SM2L30 and SM2V10, Thorlabs). Position the 1000 mm and 500 mm achromatic lenses (ACT508-500-A-ML and ACT508-1000-A-ML, Thorlabs) with a separation of 100.3 mm apart (measured **edge to edge**, not centre to centre, see Figure S4.
90. (Photo 11B-C) Construct the SM2 cage system as shown in the photo. Use an SM2-to-SM1 cage adapter (LCP6, Thorlabs) to extend the cage system and attach a right-angle kinematic mirror mount M12 (KCB1C/M, Thorlabs) at the end. Insert a cage iris (CP20D, Thorlabs) before M12.
91. Mount a threaded SM2 iris (SM2D25D, Thorlabs) in front of the SM2 cage plate (LCP34/M, Thorlabs). Remove the two alignment irises, then place the cage system into the optical path.
92. (Photo 11D) Adjust the position and height of the cage system such that the collimated beam passes through the centres of both the SM2 iris and the cage iris.
93. (Photo 11E) Place a bullseye level (LVL01, Thorlabs) on top of the cage system to ensure it is properly aligned at the correct height along the pre-determined optical axis.
94. (Photo 11F) Secure the TL2 assembly onto the SM2 cage plate. The beam should now be focused while still passing through the centre of the cage plate.
95. Swap O1 with a spare nosepiece port. Determine the axial position of TL2 by sliding the cage system along the optical axis until the exiting beam is collimated. Verify collimation using a shear interferometer (3-5 mm beam shear plate). **Critical**: After setting the axial position of TL2, confirm that the beam still passes through both the SM2 iris and the cage iris (i.e., the lateral position does not change).
96. (Photo 12A) M12 should reflect the beam 90 degrees to the right along the X-axis. Adjust the knobs on M12 and use the two-iris alignment technique to ensure the reflected beam follows a well-defined horizontal optical path.
97. (Photo 12B-E) Mount O2 on a 3D translation stage TS2 (LX30/M, Thorlabs) and position it after M12 so that the transmitted beam passes through the centre of O2’s back aperture. Use a threaded SM1 iris to verify the beam’s position. A C-mount-to-SM1 adaptor (SM1A10, Thorlabs) is required. **Critical**: (Photo 11D) Use bullseye level to check the vertical tilt of O2. The orientation of O2 should align with the optical axis. and not to be tilted in any direction.
98. To determine the axial position of O2, set the galvanometric mirror (GM) to continuously scan between −0.25 V and 2.25 V using the *Scan-Auto* mode in the LabVIEW program (see Figure S3), and place a viewing card at the output of mirror M12. Because the GM is located at a plane conjugate to the pupil planes of both O1 and O2, a stationary conjugate plane is formed during scanning (see Figure S5). The location where the scanning beam appears stationary on the card corresponds to the BFP of O2, thus indicating its precise axial position.
99. Bring O1 back into the path and the exiting beam from O2 should be collimated. Note that the transmitted beam may not be perfectly collimated but slightly diverging when exiting the objective.

### Straight configuration – O2 axial with O3 (5 hours)

The alignment of TL2 and O2 described in the previous steps brings them into proximity. Their precise positions will require fine-tuning of the remote-refocusing module in the straight configuration. This ensures correct Galvo scanning performance, proper TL2 assembly, accurate positioning of TL2 and O2 and ultimately, correct system magnification.

100. (Photo 13A-B) Mount O3 with a rotational mount (LRM1, Thorlabs) onto a 2D translation stage TS3 (LX20/M, Thorlabs). A C-mount-to-SM1 adaptor (SM1A10, Thorlabs) is required.
101. (Photo 13C) Swap O1 with a spare nosepiece port and position O3 at a 0-degree angle (i.e., head-to-head) to the optical axis of O2.
102. Adjust the position of O3 to produce an unclipped, collimated output beam. The beam should exit through the centre of O3’s aperture. Use a shear interferometer (3-5 mm beam shear plate) to verify collimation.
103. (Photo 13D) Place the 200 mm tertiary tube lens TL3 (TTL200-A, Thorlabs) after O3. Ensure the beam passes through the centre of TL3, using a threaded iris (SM2D25D, Thorlabs) for alignment.
104. (Photo 13E) Place a mirror (M13) after TL3 to redirect the beam into an alternative optical path that provides sufficient clearance for camera mounting. The beam should strike the centre of M13 without clipping its edges.
105. (Photo 13E) Position a ND filter wheel (FW2AND, Thorlabs) after M13. Set the total optical density (OD) to 6 (OD4 + OD2) to reduce beam intensity and prevent damage to the camera sensor.
106. (Photo 13E) Mount the camera at the focal plane of TL3. Fine-tune the camera position to achieve a focused laser spot at the centre of the camera’s FOV and the intensity reading reaches the maximum (Figure 3A). Use the ‘Live View’ mode in Micro-manager 2.0 to monitor intensity and focus. **Critical**: The beam shape in the FOV should appear circular and symmetric in the xy plane. When the camera is placed away from the focal plane, a symmetric Airy Disc pattern should be visible.
107. Switch off the reference laser.
108. Apply a small amount of silicone immersion oil onto O1 and mount a No.1.5 coverslip onto the sample stage.
109. Place a variable line grating test target (R1L3S6P, Thorlabs) on top of the coverslip. Use the densest grating (250 lp/mm) for magnification measurements. **Do not** change the axial position of O1. **Note**: The use of a No.1.5 coverslip prevents direct contact between the immersion oil and the test target. No additional medium is needed between the coverslip and the grating. Ensure the grating side faces downwards, in direct contact with the coverslip.
110. According to Step 21, the height of O1 should have been identified and fixed during prior alignment. When the white light source is turned on, you should see focused line grating patterns in the camera FOV (Figure 3B). **Critical:** If the grating pattern shown in the FOV is not in focus, adjust the correction collar of O1 to achieve the sharpest grating pattern. **Do not** change the axial position of O1. If the focal position lies outside the range of the O1’s correction collar, this may indicate alignment issues (see troubleshooting table).
111. Capture a snapshot of the line grating pattern and save the image locally.
112. Record the current axial position of O1 and the micrometre reading on the translation stage TS3. **Note**: This defines the central height of O1. Based on Equation 1.5, the system supports ~82 µm of refocusing range centred around this position. That is, the system maintains stable magnification (e.g., 1.4×, using a silicon oil immersion O1 and an air objective O2) within ±41 µm of the central focal plane.
113. Adjust O1’s axial position in 10 µm increments above and below the central focus. At each step, refocus the image by axially translating O3 (via TS3) until the grating becomes sharp. Capture an image at each position. Detailed instructions are provided in [73].
114. This process yields a series of grating images acquired at different focal offsets.
115. Calculate the lateral magnification of the remote-refocusing system (from O1 to O2) using grating images. First, determine the measured pixel size by dividing a known physical distance on the grating by the number of pixels spanning that distance (measured pixel size = known distance / number of pixels). Then, calculate the lateral magnification by dividing the camera pixel size by the product of the measured pixel size and the total magnification of Microscope 3 (O3 + TL3).
116. Plot the overall lateral magnification from O1 to O2 as a function of O1 defocus (Figure 3C).
117. Adjust O2 axially to obtain a relatively stable overall lateral magnification line (typically the gradient is less than 5×10^-5^ [73].
118. If the observed lateral magnification deviates significantly from the theoretical value (e.g., 1.4 based on the refractive index ratio), adjust the spacing of lenses within the TL2 assembly and repeat Steps 88–110. **Note:** The amount to adjust each time depends on how far the TL2 position is from optimal alignment. As a reference, one full turn of the Thorlabs SM2 lens tube (40 threads per inch) corresponds to a linear movement of approximately 0.635 mm.
119. Once a stable magnification close to the desired value of 1.4 is achieved across the ~82 µm refocus range, the alignment of TL2 and O2 is considered complete. The system is now capable of remote refocusing over this range.
120. Return O1 to the initial axial position.
121. The straight transmission alignment is now complete. Proceed to alignment in the tilt configuration.

### Tilt configuration (3 hours)

122. Remove the test target and coverslip.
123. Switch on the reference laser.
124. (Photo 14A-B) Since the focal length of TL2 and the axial position of O2 are well aligned, swap O1 with a spare nosepiece port and position O3 at a 35-degree angle to the optical axis of O2.
125. (Photo 14C-D) Follow Step 95 – 99 to align the system in the tilt configuration. **Critical:** Due to the oblique geometry and the tilt of objective O3, the PSF in OPM is inherently anisotropic and typically appears slightly elliptical rather than perfectly round. When the camera is placed away from the focal plane, an elliptical Airy Disc pattern should be visible.
126. For multi-colour imaging, place the optical filter wheel FW (FW2A, Thorlabs) in front of the camera.

**Figure 3:**
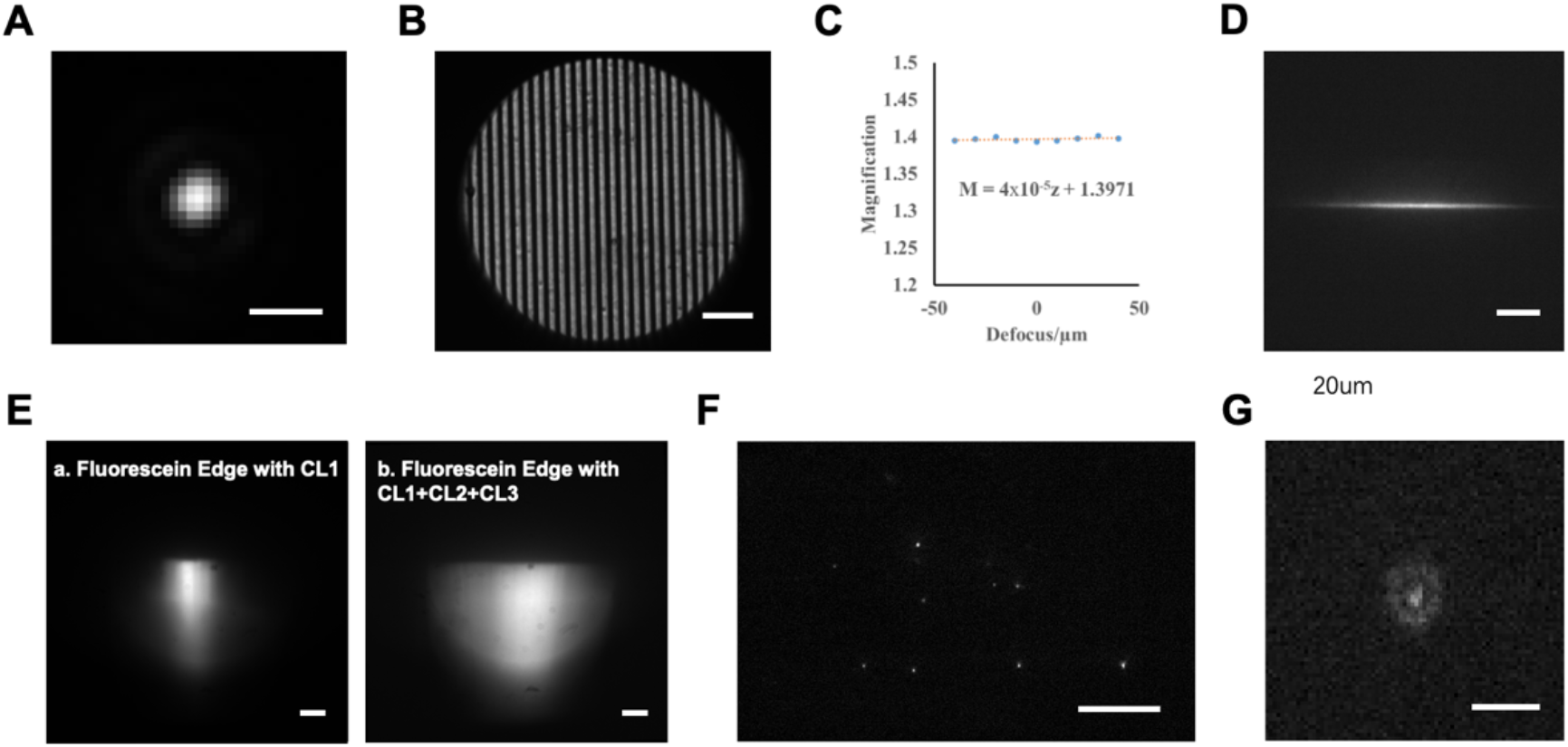
Expected results during system alignment. (A) Focused laser spot in the straight configuration. Scale bar: 1 μm. (B) Focused grating pattern in the straight configuration. Scale bar: 20 μm. (C) System magnification plotted against defocus from the central O1 position; the intercept (1.3971) indicates the measured lateral magnification from O1 to O2, closely matching the theoretical value of 1.4. (D) Focused, non-tilted light sheet in the tilted configuration. Scale bar: 20 μm. (E) Fluorescein edge images acquired without and with the additional cylindrical lens pair (CL2 and CL3). Scale bars: 20 μm. (F) Example image of fluorescence microspheres with the light sheet optimally tilted. Scale bar: 20 μm. (G) Defocused PSF in the tilt configuration, showing an elliptical pattern. Scale bar: 2 μm.

### Light sheet generation (2 hours)

127. Switch off the reference laser and bring O1 back into the path.
128. Apply a drop of silicone immersion oil if the oil is not enough. Mount a No.1.5 coverslip onto the sample stage.
129. For light sheet characterisation, a stock fluorescein solution (5 mM, 46955, Fluka) diluted at a concentration of 1:1000 in phosphate-buffered saline (PBS) is used.
130. Apply the diluted fluorescein solution onto the coverslip and excite it by the 488-nm laser.
131. Remove the ND filter wheel in front of the camera and add a 525-emission filter (MF525-39, Thorlabs) in front of the camera. The resulting fluorescence image should display a focused, horizontal light sheet in the camera FOV (Figure 3D). If the light sheet appears defocused, adjust the correction collar of O1 to achieve the most focused light sheet.
132. Acquire a snapshot of the focused light sheet and save it locally.
133. The thickness and the width of the light-sheet is measured by 1D Gaussian fitting of the FWHMs along the x and scanning direction y of the sheet profile. The corresponding Rayleigh length of the sheet can be calculated according to Equation (1.2) with the measured thickness.
134. Continue from Step 79, fine-tune the lateral position of the cylindrical lens CL1 by rotating it 360 degrees. When viewed through the camera, the light sheet should rotate around a central point. Do not alter the axial position of CL1.
135. Use the diluted fluorescein solution (5 µM, 46955, Fluka) to assist in the alignment of the additional cylindrical lens pair (CL2 and CL3; ACY254-050-A and ACY254-200-A, Thorlabs), which extend the lateral light-sheet width by approximately 4-fold along the X-axis.
136. Tilt the light sheet by adjusting the translation stage TS1 along the X-direction to reposition the dichroic mirror DM2.
137. You should see a sharp fluorescein edge at the centre of the field of view (see Figure 6). This will serve as your reference for aligning the cylindrical lens pair. Note that moving O1 away from the optimal focus position will cause the edge to appear blurred, indicating defocus.
138. Insert cylindrical lens CL2 between CL1 and mirror M6, ensuring that the curved surface of CL2 faces CL1.
139. Align the rotation mount of CL2 to match the rotational orientation of CL1.
140. Fine-tune the height and lateral position of CL2 until the sharp fluorescein edge is restored at the centre of the field of view.
141. Insert cylindrical lens CL3 between mirror M6 and dichroic mirror DM1.
142. Align the rotation mount of CL3 to match those of CL1 and CL2.
143. Adjust the height and lateral position of CL3 to again produce a sharp fluorescein edge at the centre of the FOV.
144. Finally, adjust the axial position of CL3 to identify the position where the fluorescein edge appears widest, indicating optimal alignment of the expanded light sheet (Figure 3E).
145. The entire system alignment is now complete. Proceed to system characterisation.

### System characterisation with fluorescence microspheres (3 hours)

146. Prepare a sample of fluorescent microspheres embedded in agarose on a No.1.5 coverslip. See SI for detailed sample preparation protocols.
147. Remove the fluorescein sample and mount the agarose microsphere sample onto the sample stage.
148. The fluorescent microspheres should appear in focus in the camera FOV. If they appear defocused, adjust the correction collar of O1 to obtain the most focused and symmetric PSFs in 3D.
149. With the light sheet in a non-tilted configuration, a **single line** of fluorescent microspheres should be visible in the camera FOV.
150. Translate the 1D stage TS1 to tilt the light sheet exiting O1 to approximately 35 degrees relative to the horizontal plane.
151. Fine-tune the tilt angle of the light sheet while monitoring the image to ensure:
  - Fluorescent microspheres remain in focus across a stripe in the FOV, rather than a narrow line (Figure 3F).
  - The system achieves the highest signal-to-noise ratio (SNR).
  - The PSFs of the microspheres appear elliptical without clipping or distortion when defocused (Figure 3G). **Critical:** Any signs of PSF clipping or distortion may indicate misalignment of O3. See troubleshooting table for more details.
152. When adjusting the axial focus of O1, microspheres located in the central region of the FOV, where the light sheet is thinnest and most optimised, should not appear defocused. Instead, they should “blink”, meaning they appear and disappear as sharp points without significant defocusing.
153. When translating the sample stage along the scanning direction, a similar blinking behaviour should be observed, consistent with step 151.
154. Use the LabView control and Micro-Manager script to scan the galvo mirror and acquire a volumetric image stack of the fluorescent microspheres. Refer to SI for detailed acquisition instructions.
155. 3D reconstruction of the volumetric image stack can be achieved by a custom python analysis script. Refer to SI for detailed explanation and instructions.

### Anticipated results

Successful implementation of the protocol produces a single-objective OPM system capable of achieving near-diffraction-limited resolution, with an average lateral (xy) resolution of approximately 374 nm and an axial (z) resolution of 660 nm, as measured across ten 100 nm diameter fluorescent microspheres with an excitation wavelength of 488 nm. A three-dimensional maximum intensity projection of the bead sample is shown in Figure 4A. The system’s resolution was quantified by measuring the full width at half maximum (FWHM) of the intensity profiles for individual fluorescent microsphere images. An example bead is highlighted in figure 4B, with representative lateral (xy) and axial (xz) intensity profiles and their corresponding Gaussian fits displayed in Figure 4C. These results demonstrate that the constructed OPM setup achieves the expected optical performance for three-dimensional fluorescence imaging at submicron resolution.

**Figure 4:**
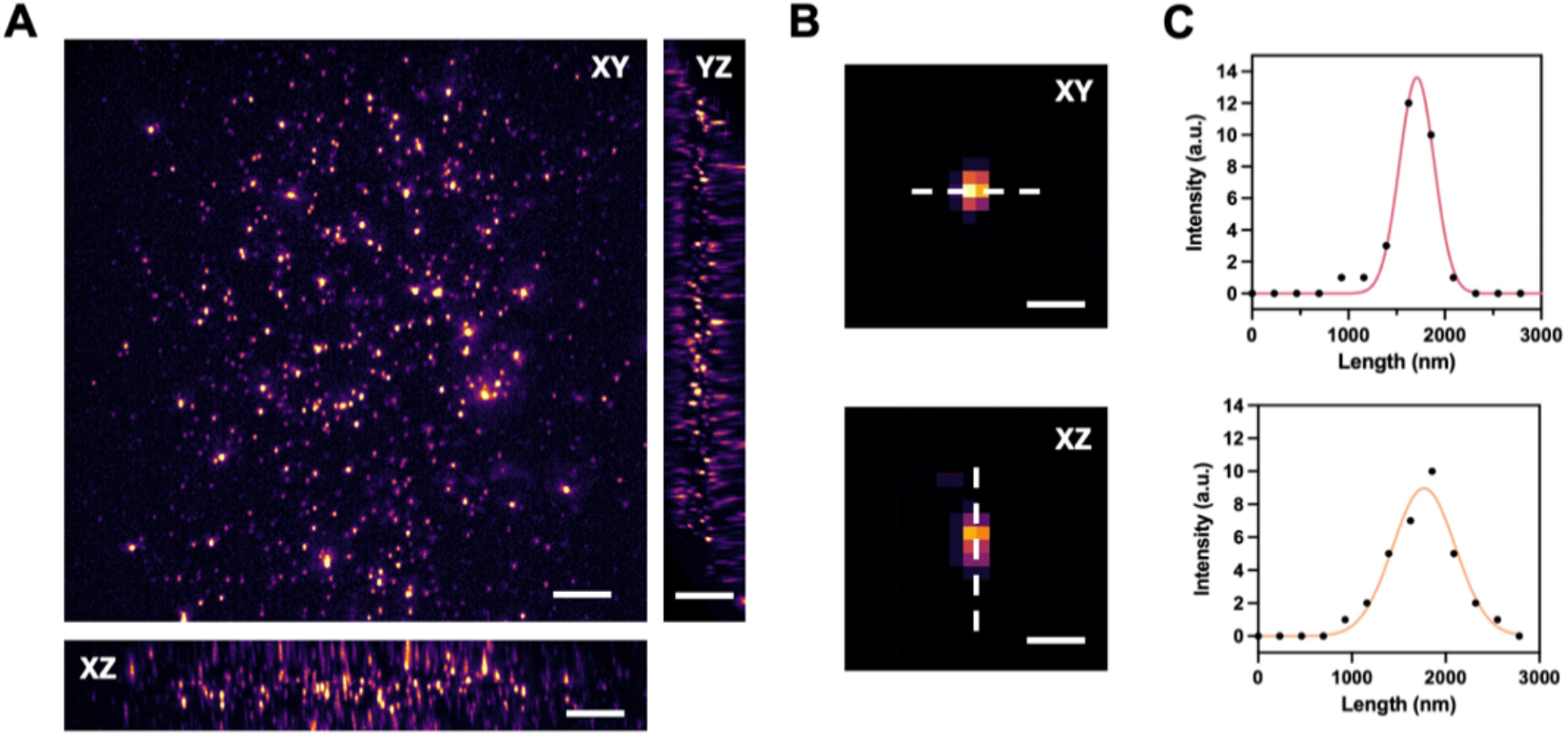
System resolution characterisation using sub-diffraction fluorescent microspheres. (A) Three-dimensional maximum intensity projections of fluorescent microspheres acquired with the single-objective OPM system, showing the XY, XZ, and YZ planes. Scale bars: 10 μm. (B) Representative bead image displayed in the XY and XZ planes, highlighting the regions used for intensity profile analysis (dashed lines). Scale bars: 1 μm. (C) Lateral (top) and axial (bottom) intensity profiles through the bead, with corresponding Gaussian fits (coloured lines), used to determine the full width at half maximum (FWHM) and quantify system resolution.

**Figure 5:**
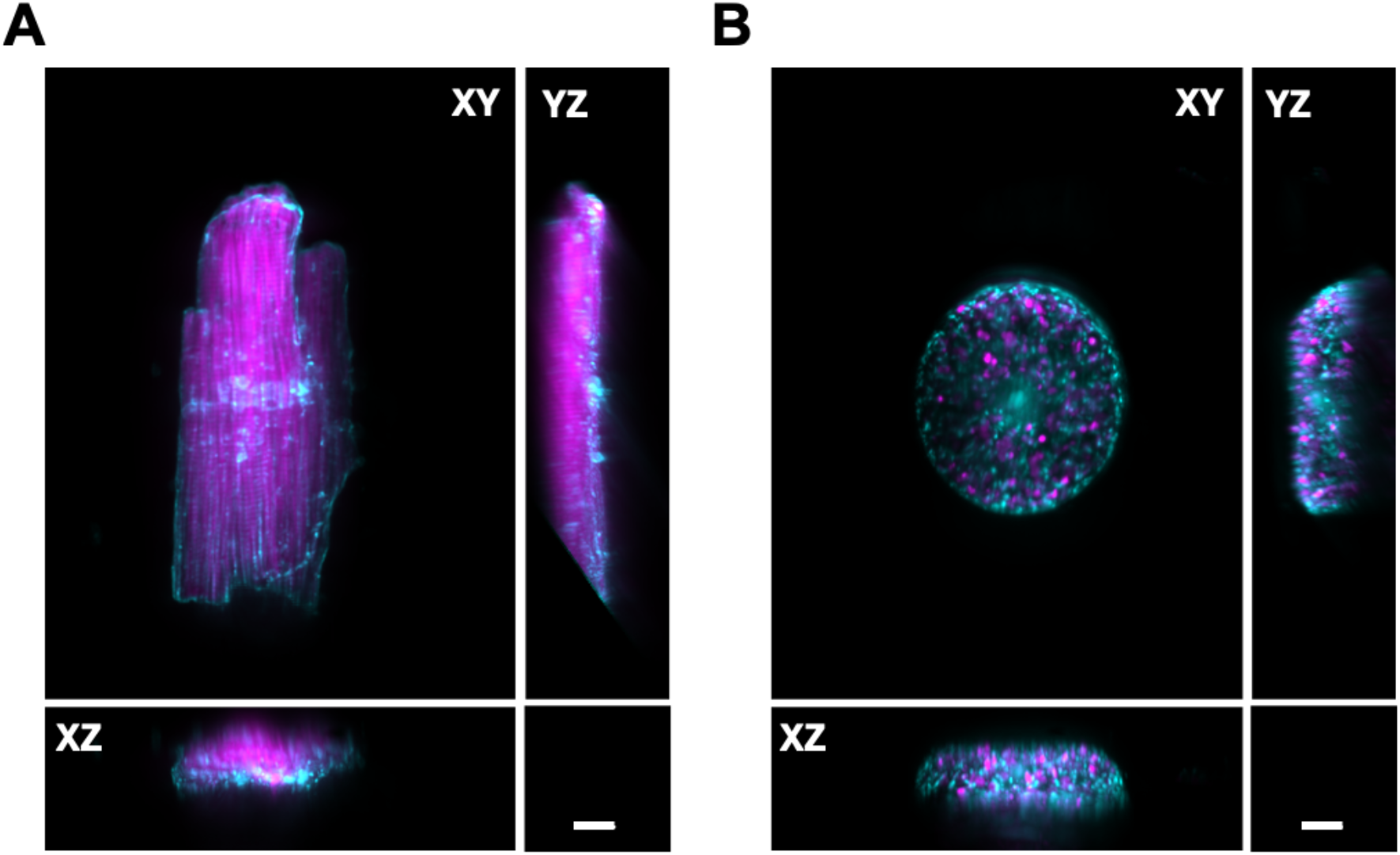
High-resolution two-colour imaging of cardiomyocytes and diatoms using the single-objective OPM. (A) Representative volumetric reconstructions of adult rat ventricular cardiomyocytes acquired with the single-objective OPM. Cardiomyocytes were stained with WGA–Alexa Fluor™ 488 (cyan) to label cell surface glycoconjugates, highlighting membrane structures. CellMask™ Deep Red (magenta) was used to stain the actin cytoskeleton, revealing the highly ordered myofibrillar architecture of the cardiomyocytes. (B) Representative volumetric reconstructions of *Coscinodiscus radiatus* diatoms. Diatoms (Coscinodiscus radiatus) were labelled with SYTO® 9 (cyan) to visualise nucleic acids, marking the nucleus and other DNA/RNA-rich regions. Chloroplast autofluorescence is shown in magenta, outlining the internal organelle structure. XY and XZ (YZ) views illustrate the system’s ability to resolve 3D structural with high-resolution across morphologically distinct specimens. Scale bars: 10 µm.

To showcase the capabilities of the single-objective OPM system, we applied it to two biologically and structurally distinct sample types: mammalian cardiomyocytes and marine diatoms. Adult rat ventricular cardiomyocytes (ARVMs) were isolated, cultured, and fluorescently labelled with WGA–Alexa Fluor™ 488 to visualise cell surface glycoconjugates and membrane structures, and with CellMask™ Deep Red to stain the actin cytoskeleton.

Volumetric reconstructions revealed the highly ordered myofibrillar architecture and detailed membrane morphology in both top-down and side views. In parallel, the diatom Coscinodiscus radiatus was used to demonstrate the system’s ability to resolve three-dimensional intracellular architecture within rigid silica frustules.

Diatoms were labelled with SYTO® 9 to visualise nucleic acids, while chlorophyll autofluorescence provided intrinsic contrast for internal plastid structures. These examples illustrate the system’s ability to deliver high-resolution, multi-colour volumetric imaging across morphologically distinct specimens, capturing fine structural details in diverse biological contexts.

## Supporting information

Alignment Guide

OPM_Easy2Use_Form

Partlist

## Code availability

Complete-suite-for-single-objective-OPM: https://github.com/Zui409/Complete-suite-for-single-objective-OPM

## Author contributions

The manuscript was written by ZZ, WH, and YW. The system was designed by ZZ and WH. The system construction and characterisation were conducted by ZZ, WH and AD. The LabVIEW control program was written by WH, and the image reconstruction scripts were developed by ZZ and YW. Imaging experiments were conducted by ZZ, WH and AS. ZZ, AD annotated the photos in the alignment procedure. AS and DK supervised the entire project. All authors reviewed the manuscript.

## Acknowledgement

The authors gratefully acknowledge Dr Fengjie Liu for providing the diatom samples and Professor Julia Gorelik for providing the adult rat cardiomyocyte samples used for the system demonstration.

## Troubleshooting table

Troubleshooting table:
| Step | Problem | Possible Reason | Solution |
| --- | --- | --- | --- |
| 7 | The reference beam shifts significantly after inserting or removing O1. | The optical axis defined by the two irises is not perfectly vertical. | Ensure the two irises are mounted securely when aligning the reference laser. Ensure a long separation between them to improve accuracy, consider using a longer lens tube if space allows. Carefully fine-tune the kinematic collimator adaptor until the beam passes through the centre of both irises. This step has minimal tolerance for error. |
| 60-67 | The beam passes through the alignment irises but drifts off-axis after inserting lenses L3 and L4. | The cage system may be slightly twisted or misaligned with the optical axis. L3 and L4 may not be centred within the cage system so that the beam may not pass through the centres of both lenses. | Slightly adjust the height and position of the cage system so the beam passes through the alignment irises with the lenses inserted. Any residual offset can be corrected using downstream mirrors M3 and M4. |
| 72, 73 | The reference beam and the excitation beam do not overlap. | Typically caused by one or both beams not passing through the centres of the lenses. | Determine whether the reference or excitation beam is misaligned. Check that the reference beam passes through the centres of L4 to L7. Use alignment irises at multiple positions to verify beam centring. Avoid adjusting optics in the common path unless necessary, as it affects emission path alignment. If the reference beam is aligned properly and the problem persists, fine-tune M3 and M4 to match the excitation beam to the reference. |
| 77 | The beam exiting O1 is not collimated. | The three 4f systems (O1–TL1, L7–L6, and L5–L9 (CL1)) are not properly aligned. | Use a shear interferometer to verify collimation through each 4f system. Realign if necessary. |
|  |  | The beam expander may be incorrectly spaced. | Adjust the distance between L3 and L4 in the beam expander cage system until minimal divergence is achieved at the output of O1. |
| 85 | When aligning additional laser lines, the beams do not overlap with the reference beam or with each other | Misalignment in the beam expander cage system or the mirrors in the excitation path. | Verify that the laser passes through the centres of the beam expander lenses. Realign if necessary. Then adjust the periscope mirrors (e.g., M1 and M2 for the 638-nm laser) to match the excitation beams to the reference beam. |
| 98 | O1 focus shifts along the optical axis during galvo scanning. | The galvo mirror is not correctly positioned at the plane conjugate to the pupil planes of O1 and O2. | Carefully readjust the galvo mirror position to ensure proper conjugation, then realign the system starting from Step 27. |
| 106 | Beam shape is not symmetrical in xy. | Misalignment of O2 and O3. | Repeat Step 100 – 106 to realign the remote-refocusing module. |
|  |  | Clipping at mirrors or irises. | Clipping may occur at any mirror or closed iris in the emission path.<br>Ensure all irises are fully open.<br>Inspect for beam clipping at all mirrors, and if detected, reposition the affected mirror and realign all downstream optics accordingly. |
| 109 | Grating pattern is not visible in the camera FOV. | Grating side is facing upwards and not in contact with the coverslip. | Flip the grating so the patterned side faces downwards in direct contact with the coverslip. |
|  |  | Coverslip thickness is not appropriate. | Use a No. 1.5 glass-bottom coverslip to ensure proper optical path length. |
|  |  | Insufficient immersion oil on O1. | Add one or two drops of Silicone immersion oil to O1 to ensure full contact. |
| 110 | Grating pattern in the camera FOV is not in focus. | O1 correction collar not correctly set. | Adjust the correction collar on O1 to sharpen the grating pattern. **Do not** change the axial position of O1. |
|  |  | If the focal plane is outside the collar range, the image plane of O1 is not conjugated with the camera image plane. One or more 4f systems (O1–TL1, L7–L8, TL2–O2, O3–TL3) may be misaligned. | Set the objector collar to 0.17 mm, which corresponds to the position for a No.1.5 coverslip.<br>Verify alignment of the reference beam through each 4f system using a shear interferometer.<br>If L7–L8, TL2–O2, and O3–TL3 are correctly aligned, then adjust the axial distance between O1 and TL1.<br>Refocus O1 on the grating and fix it at that height for subsequent alignment. |
| 115 | Incorrect lateral magnification from O1 to O2. | TL2 has an incorrect focal length.<br>Incorrect separation between the two doublet lenses. | Adjust the separation between the two doublets within the SM2 lens tube. If the measured lateral magnification is greater than the expected value, increase the lens separation accordingly, and vice versa. After adjustment, realign the system starting from Step 91. |
| 125 | Beam shape is not elliptical or appears distorted. | Misalignment of O2 and O3. | Repeat Steps 97 – 103 to realign the remote-refocusing module until a clean elliptical beam shape is observed in the camera FOV. |
|  |  | Clipping at mirrors or irises. | Clipping may occur at any mirror or closed iris in the emission path.<br>Ensure all irises are fully open.<br>Inspect for beam clipping at all mirrors, and if detected, reposition the affected mirror and realign all downstream optics accordingly. |
| 131 | The light sheet in the camera FOV is not in focus. | O1 correction collar not correctly set. | Adjust the correction collar on O1 to achieve the sharpest light sheet. **Do not** change the axial position of O1. |
|  |  | The axial position of the cylindrical lens CL1 is incorrect. | Set the objector collar to 0.17 mm, which corresponds to the position for a No.1.5 coverslip.<br>Avoid altering the axial position of CL1, as it is shared between the 488-nm and 638-nm channels.<br>Instead, slightly adjust the position of L4 in the beam expander cage system until the light sheet appears sharpest in the camera FOV. |
| 131 | The light sheet in the camera FOV is not horizontal. | Incorrect rotation angle of CL1. | Rotate CL1 until a horizontal light sheet is visible in the camera FOV. |
| 151 | The PSF of the fluorescent microspheres is not elliptical or appears distorted. | Misalignment of O2 and O3. | Repeat Steps 97 – 103 to realign the remote-refocusing module until unclipped elliptical PSFs are visible in the FOV. |
| 151 | The fluorescent microspheres in the camera FOV are not in focus. | O1 correction collar not correctly set. | Adjust the correction collar of O1 to achieve the focused fluorescent microspheres. **Do not** change the axial position of O1. |
|  |  | The axial position of O3 is slightly off. | Set the objector collar to 0.17 mm, which corresponds to the position for a No.1.5 coverslip.<br>Slowly translate O3 along the axial direction using the translation stage TS3 until the majority of the fluorescent microspheres appear in focus in the camera FOV. |

