## Supplementary material for "Constructing a single-objective oblique plane microscope (OPM) for fast, multi-colour, high-resolution volumetric fluorescence imaging": Alignment Guide

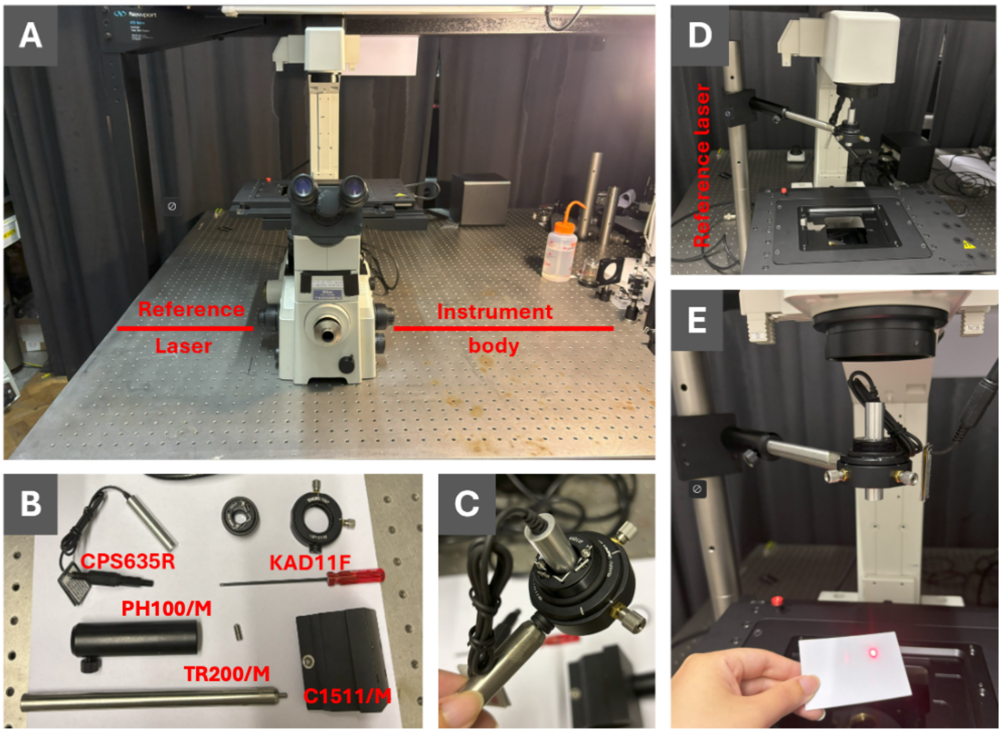


**Photo 1: Reference path setup for microscope alignment. (Step 1-2)**
(A) Positioning the microscope body on a stable optical table, with approximately 15 cm reserved on the left for the reference laser assembly and 80 cm on the right for construction of the main instrument. (B) Components for assembling the reference laser path, including the collimated reference laser (CPS635R), kinematic collimator adaptor (KAD11F), 200 mm post (TR200/M), 100 mm post holder (PH100/M), and post mounting clamp (C1511/M). (C) Installation of the collimated reference laser onto the kinematic adaptor for precise adjustment of beam position and tilt. (D) The reference laser and adaptor are mounted around 15 cm above the sample stage on a 50 cm tall, 1.5″ diameter vertical post. (E) Final configuration, with the collimated reference beam propagating vertically downward toward the microscope nosepiece, ready for optical axis alignment.

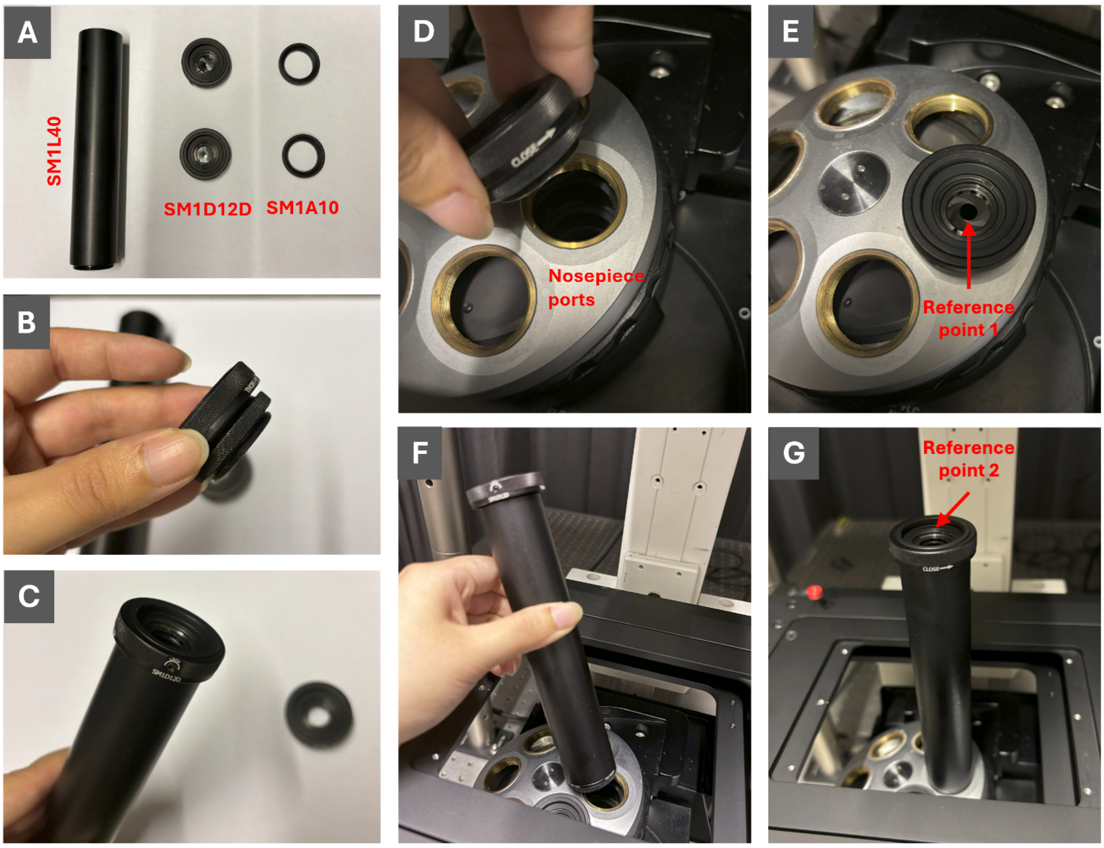


**Photo 2: Defining the vertical optical axis for microscope alignment. (Step 3-4)**
(A) Components for iris mounting: 4.00″ lens tube (SM1L40), threaded iris (SM1D12D), and SM1-to-C-mount adapter (SM1A10). (B, C) Assembly of the threaded iris with the adapter, and with the lens tube. (D, E) Installation of the threaded iris (with adapter) onto one nosepiece port of the microscope, establishing the first reference point (Reference point 1). (F, G) Installation of the second threaded iris, assembled with the 4.00″ lens tube and adapter, onto an adjacent nosepiece port, creating the second reference point (Reference point 2). The centres of these two irises define a precise vertical optical axis for all downstream alignments.

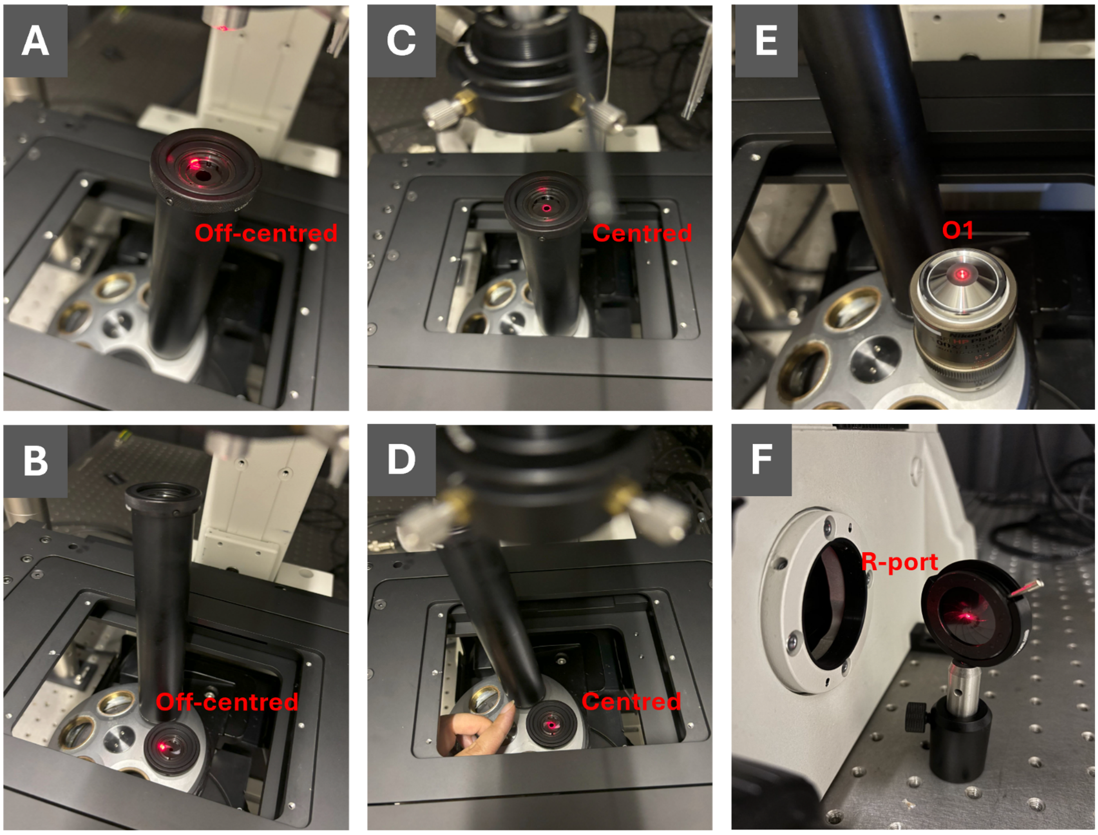


**Photo 3: Aligning the reference laser to define the vertical optical axis. (Step 5-9)**
(A, B) The reference laser beam is initially off centred with respect to the irises. (C, D) Adjustment of the collimator adaptor allows the reference beam to pass through the centres of both irises, establishing a vertically aligned optical axis. (E) The aligned reference beam is centred on the objective lens (O1). (F) The focused reference beam is visible through the R-port of the microscope body with the O1 removed from the optical path.

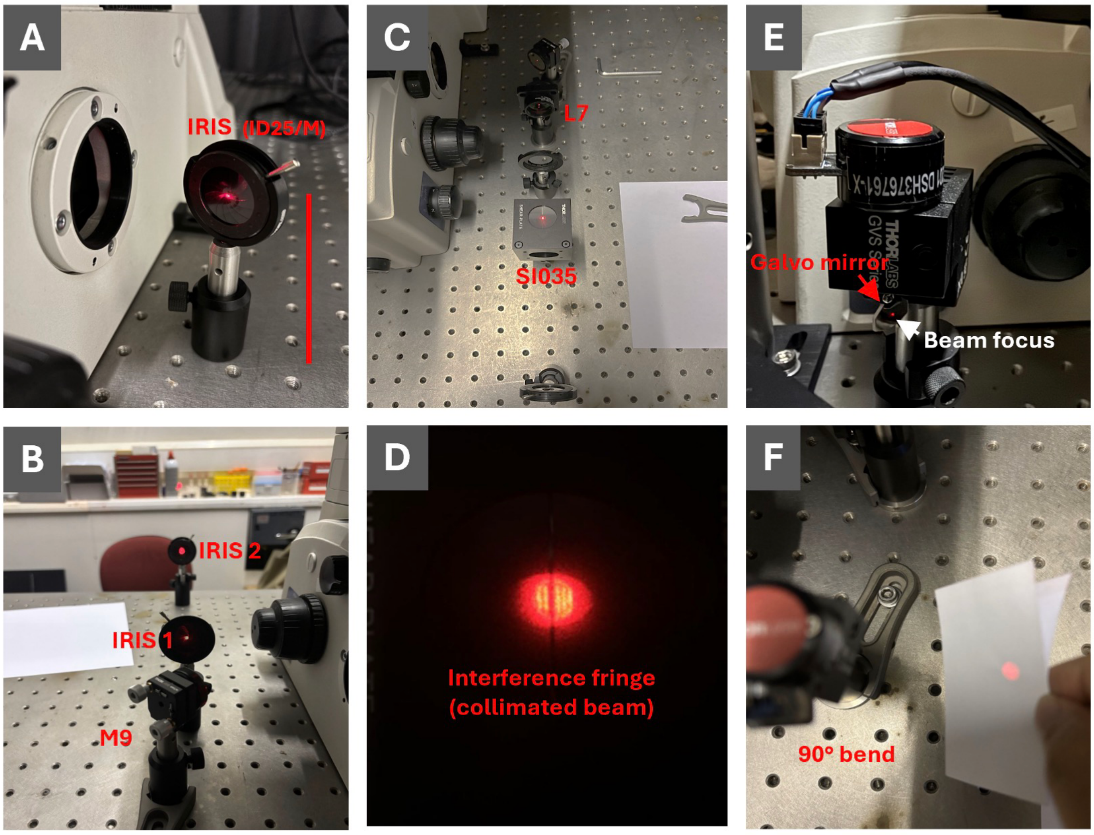


**Photo 4: Aligning the common optical path at the R-port of the microscope. (Step 12-17, 27)** (A) Determining the height of the optical axis emerging from the R-port using a post-mounted iris (ID25/M). The aperture height is adjusted so that the reference beam passes centrally through the iris. (B) Positioning a second iris and a mirror (M9) near the focal point of the reference beam; M9 reflects the beam by 90°, and the two-iris alignment technique is used to ensure the reflected beam passes through both iris centres, defining a horizontal optical axis. (C) Mounting a threaded iris in front of the achromatic lens (L7) and aligning the lens along the optical path so that the beam passes centrally through the lens and emerges collimated. (D) Verification of beam collimation using a shear interferometer (SI035) and magnifier (SIVS). Parallel interference fringes indicate a well-collimated beam. (E) Setup of the galvo mirror (GM) at a fixed voltage (1.5 V), with the beam sharply focused at its centre. (F) Observation of the reflected beam as it diverges (defocuses) along the optical axis after the 90° bend.

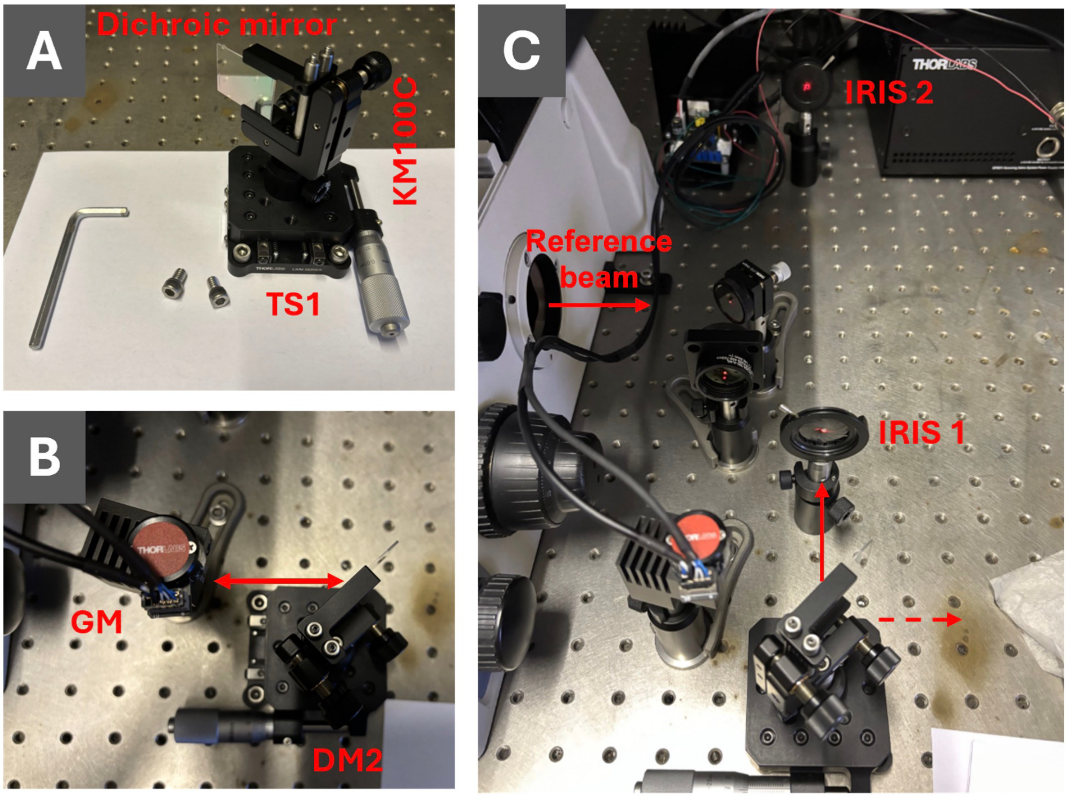


**Photo 5: Setting up the dichroic mirror (DM2) on a translation stage. (Step 29-32)**

(A) Quad-band dichroic mirror (DM2) secured in a rectangular optics holder (KM100C) and mounted on a one-dimensional translation stage (TS1). (B) TS1 is oriented to move along the X-axis of the optical table, with DM2 positioned approximately 5 cm from the galvo mirror (GM) to allow space for downstream optics. (C) The dichroic mirror (DM2) reflects the reference beam by 90° along the Y-axis (solid arrow). The Two-Iris Alignment Technique is used to ensure precise alignment of the reflected path. A dimly transmitted beam continues along the X-axis (dashed arrow), which will later be used for aligning the emission path.

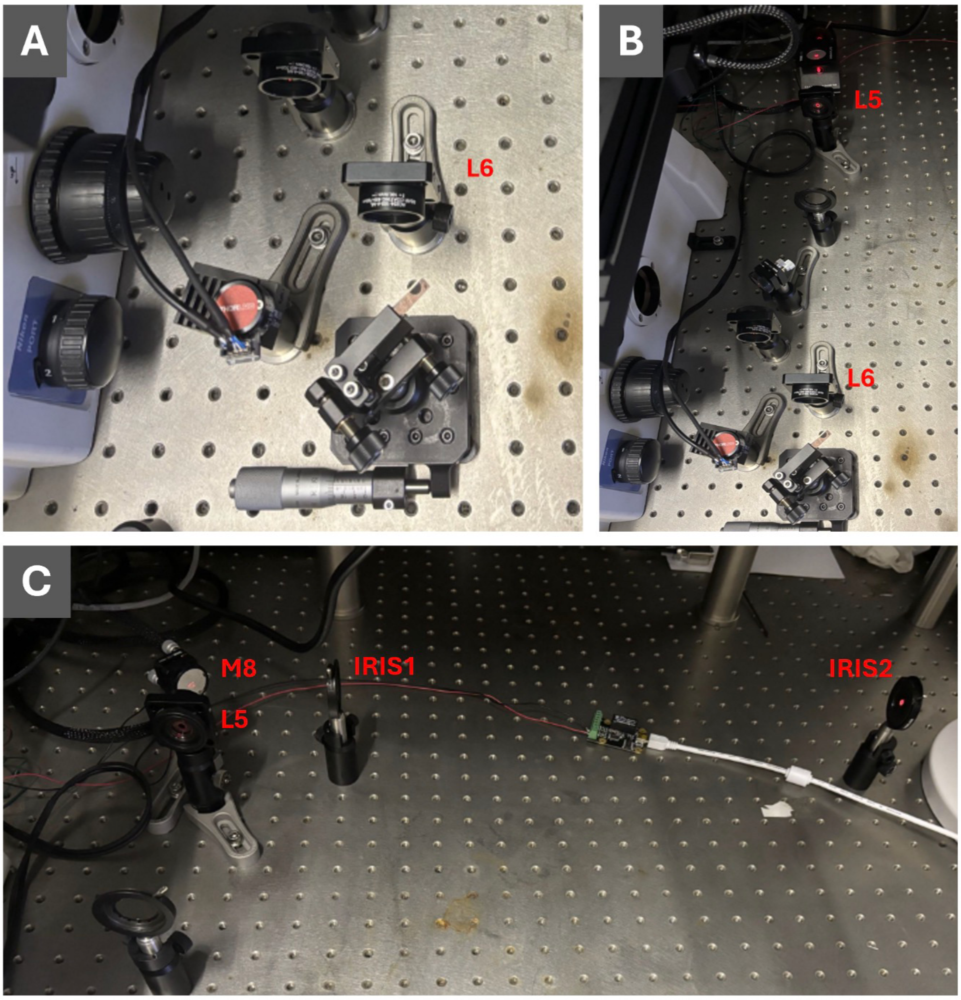


**Photo 6: Setting up the excitation path. (Step 37-40)**

(A) With objective O1 in the optical path, a 100 mm achromatic lens (L6) is positioned such that the reference beam exiting L6 is collimated. (B) After removing O1 from the system, a 300 mm achromatic lens (L5) is positioned so that the beam exiting L5 remains collimated. Proper positioning is confirmed by beam collimation checks using a shear interferometer before and after L6 and L5, respectively. (C) A mirror (M8) is placed immediately after L5 to reflect the collimated beam by 90° to the right. The Two-Iris Alignment Technique is used to align the beam along the new direction.

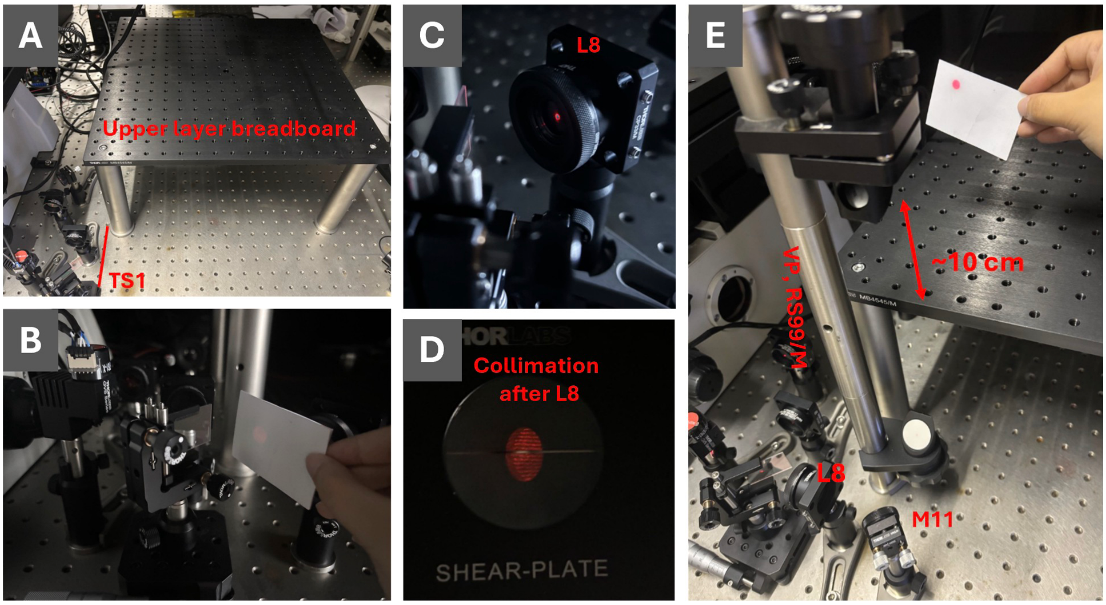


**Photo 7: Approximate positioning and alignment of the upper layer. (Step 41-46)**

(A) Three 1.5” diameter, 200 mm posts are used to support the upper breadboard layer, with the left edge of the breadboard aligned to the right edge of the 1D translation stage (TS1) as indicated by the red line. The upper right corner (hidden in the photo) is left unsupported to allow for laser mounting beneath. (B) Detection of the weak transmitted reference beam through the dichroic mirror (DM2) in the dark, with O1 in the optical path and the beam focused at the galvo mirror.

(C) Positioning of lens L8 (AC254-100-A-ML) using the weak transmitted reference beam for alignment. (D) Checking collimation of the beam after L8 with a shear interferometer (1–3 mm shear plate), with O1 in the optical path. (E) Mirror M11 is placed approximately 5 cm after L8 to reflect the beam by 90°, and the vertical periscope (VP) system (RS99/M) is assembled and positioned to direct the beam upward to the upper layer. The optical axis for the upper layer is set approximately 10 cm above the breadboard.

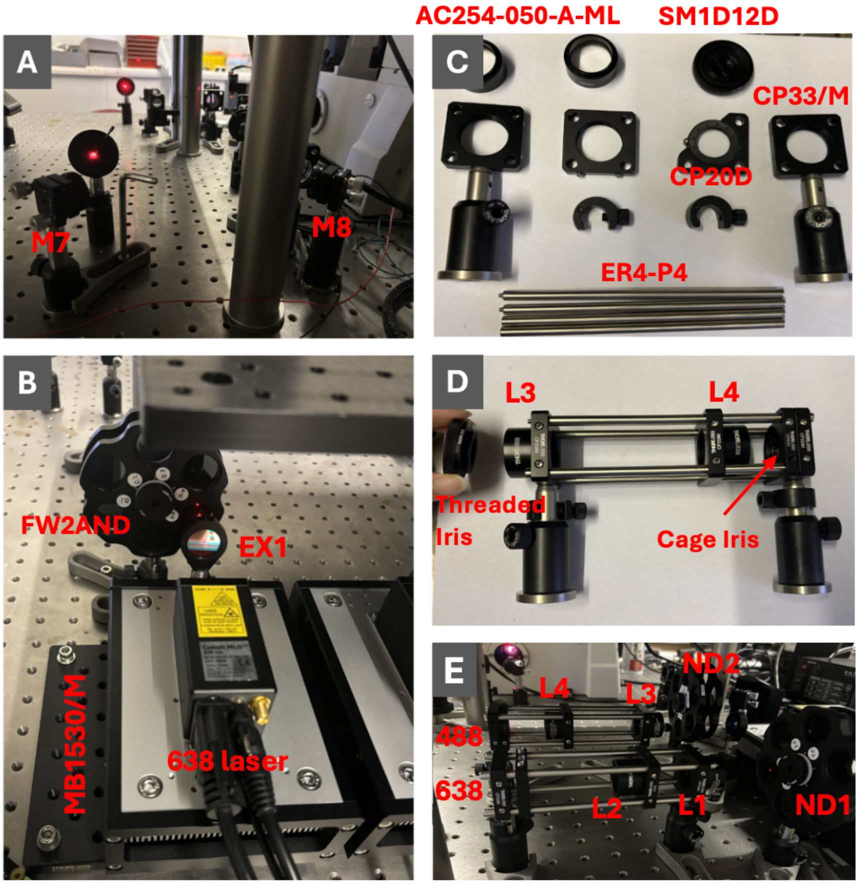


**Photo 8: Alignment of the excitation path. (Step 54-65)**

(A) Placement of mirror M7 to reflect the beam by 90° along the Y-axis, ensuring the beam does not clip at M11 and the vertical periscope. (B) Alignment of the excitation path from the laser head. Lasers are mounted with heatsinks on a breadboard (MB1530/M) beneath the upper layer, ensuring the output beam height matches the pre-defined optical axis. The neutral density filter wheel (FW2AND) and excitation filter (EX1) are placed sequentially along the path. (C) Assembly of the SM1 cage system with cage plates (CP33/M), cage rods (ER4-P4), and initial alignment using irises to define the optical axis. (D) Assembly of the beam expander lenses (L3 and L4, both AC254-050-A-ML) and a cage iris (CP20D) into the cage system, with a threaded iris mounted on the front cage plate for alignment. (E) Integration of the assembled beam expander cage systems, after the ND filters, into the excitation paths for both 488 nm and 638 nm channels.

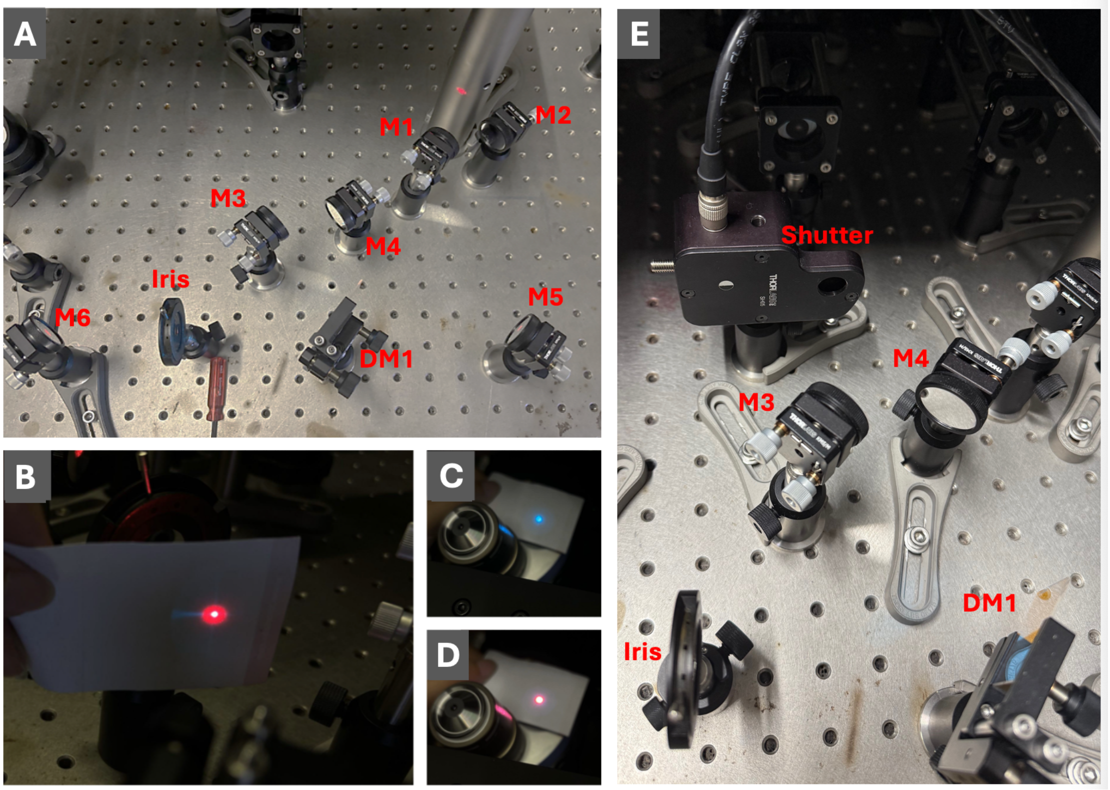


**Photo 9: Alignment of the excitation path continued. (Step 69-74)**
(A) Positioning mirrors M3 & M4 (for the 488 nm channel), and M1 & M2 (for the 638 nm channel) sequentially after the beam expander. M3 and M4 form a periscope system for fine adjustment of the 488 nm beam direction. Place M5 after M1 and M2 in the 638 nm channel to reflect the beam towards the microscopy body. The single-band dichroic mirror DM1, mounted on a kinematic mount, combines the 488 nm and 638 nm excitation beams by reflecting the 488 nm beam. (B) Overlap check of the excitation and reference beams using an alignment card. (C, D) Full overlap between the 488 nm and 638 nm beams at the back focal plane of O1, as viewed on an alignment card. (E) Placement of the optical shutter (SH05R/M) in the after the beam expander in the 488 nm channel.

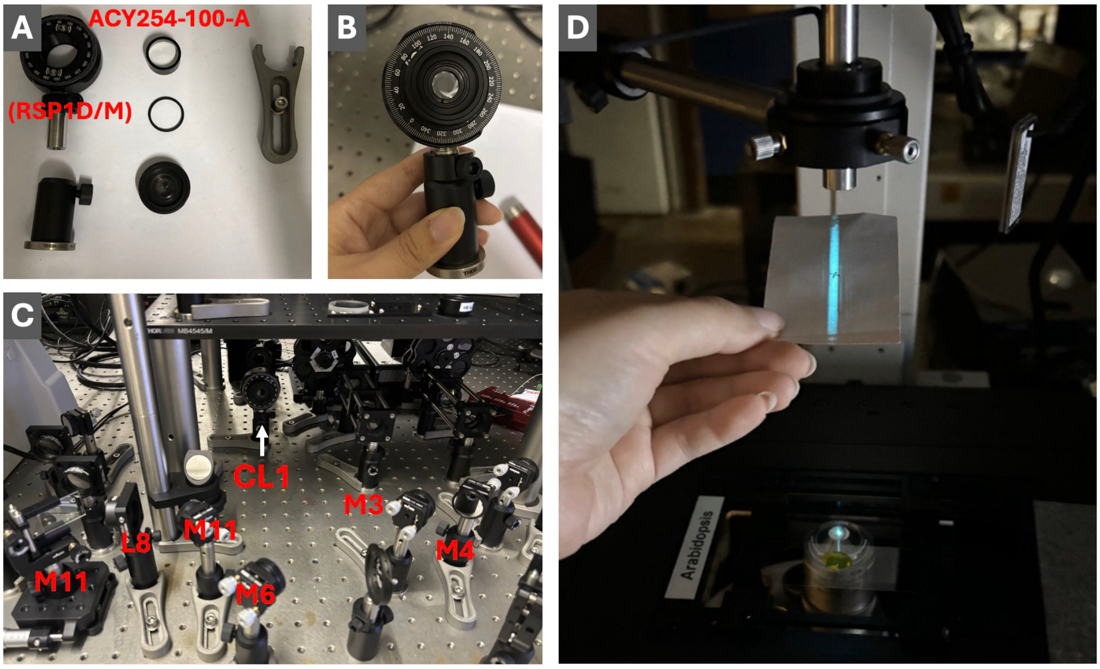


**Photo 10: Alignment of the cylindrical lens for light sheet generation. (Step 79-81)**
(A, B) Mounting the cylindrical lens (CL1, ACY254-100-A) on a rotational mount (RSP1D/M), with a threaded iris used to help identify the optical centre. (C) Placement of CL1 into the excitation path, replacing lens L9. The curved surface of the lens faces the incoming excitation beam from the laser head. (D) Adjusting the rotation and axial position along the optical axis of CL1 until a sharp light sheet (focused line along the Y-axis) is observed above the O1 objective.

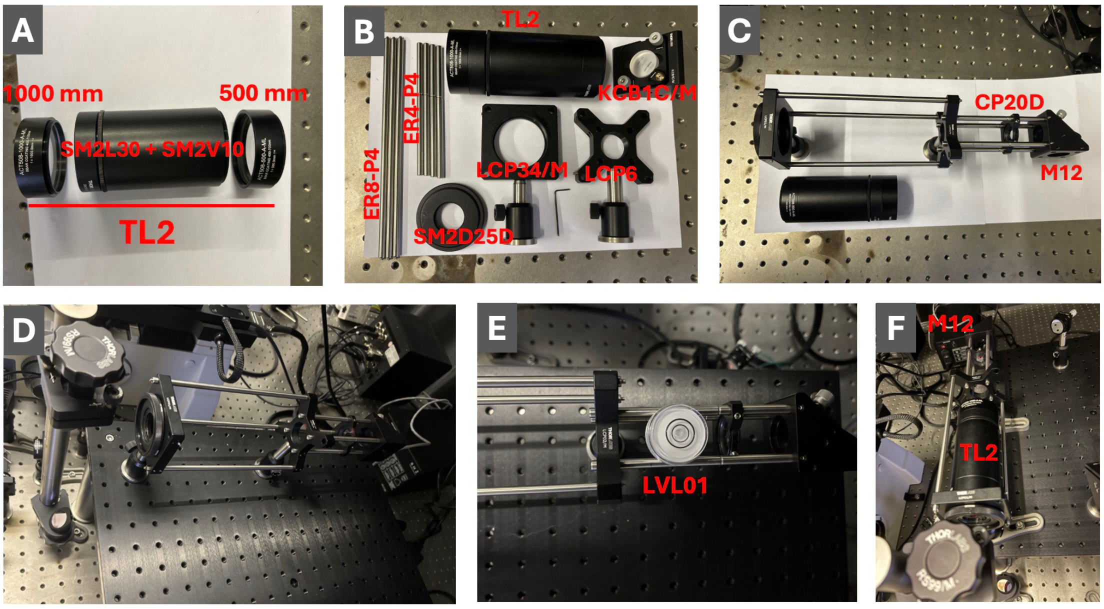


**Photo 11: Alignment and assembly of the emission path and secondary tube lens (TL2). (Step 89-94)**

(A) Assembly of the secondary tube lens (TL2) using an adjustable lens tube (SM2L30 and SM2V10, Thorlabs) with 1000 mm and 500 mm achromatic lenses, spaced 100.3 mm lens edge-to-edge. (B) Components required for constructing the TL2 cage system, including cage rods (ER8-P4 and ER4-P4), SM2 threaded iris (SM2D25D), SM2 cage plate (LCP34/M), SM2-to-SM1 cage adapter (LCP6) and right-angle kinematic mirror mount (KCB1C/M) for M12. (C) Construction of the cage system, with the extension and insertion of a cage iris (CP20D) before the mirror mount. (D) Alignment of the emission beam into the cage system using a threaded SM2 iris, ensuring the collimated beam passes through the centres of both the SM2 iris and the cage iris. (E) Placement of the assembled cage system onto the upper layer. Level check is performed using a bullseye level (LVL01) to ensure the entire TL2 is not tilted with respect to the optical axis. (F) Mounting TL2 into the cage system. The emission beam should now be focused and continue to pass centrally through both irises.

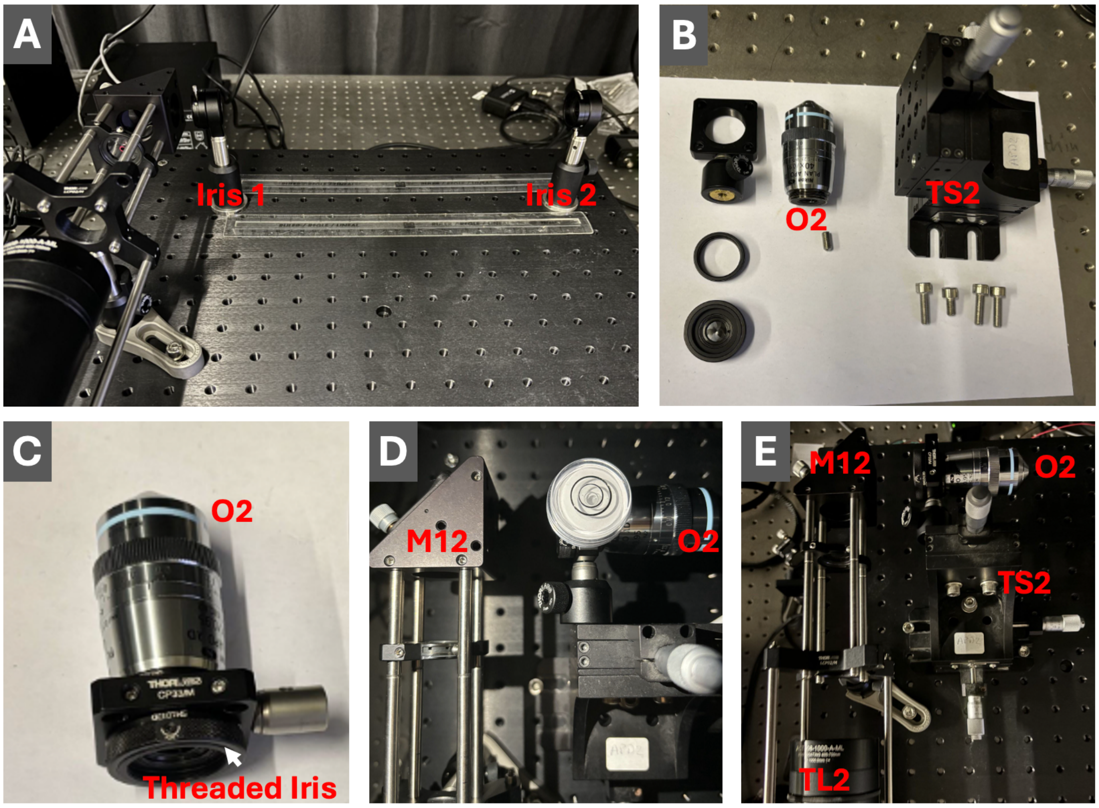


**Photo 12: Setting up and aligning the secondary objective (O2) on the upper layer. (Step 96-97)**

(A) Alignment of the horizontal optical path using two irises, with the reflected beam from mirror M12 precisely directed along the desired axis. (B, C) Mounting objective O2 on a three-dimensional translation stage (TS2, LX30/M), using a C-mount-to-SM1 adapter (SM1A10) for compatibility. A threaded SM1 iris is used to verify the beam's passage through the centre of the objective's back aperture. (D) Checking the level of TS2 using a bullseye level (LVL01) to ensure correct alignment tilt. (E) Overview of the completed optical assembly, showing TL2, M12, O2, and TS2 positioned and aligned on the upper breadboard.

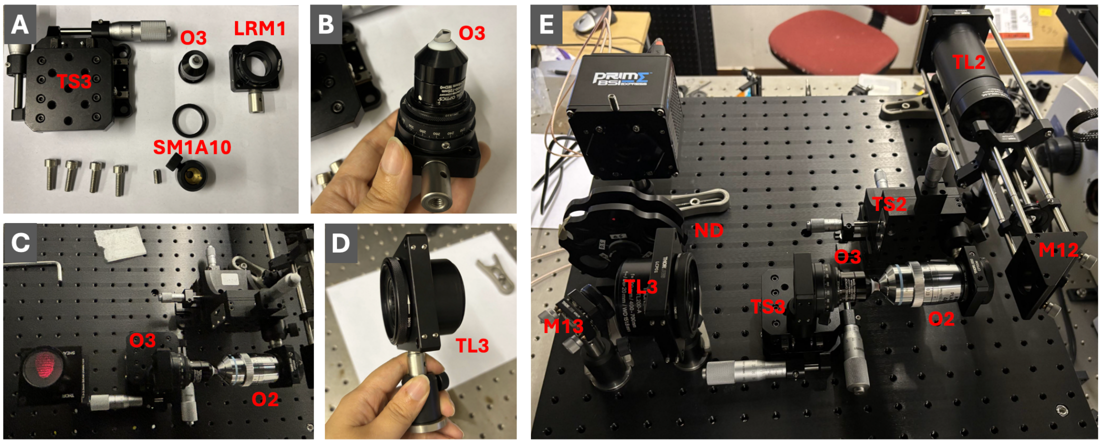


**Photo 13: Alignment of the tertiary objective (O3) and detection optics in straight configuration. (Step 100-106)**

(A, B) Components and assembly steps for mounting O3 onto a two-dimensional translation stage (TS3, LX20/M) using a rotational mount (LRM1) and a C-mount-to-SM1 adaptor (SM1A10). (C) Positioning O3 at a 0-degree angle (head-to-head) relative to the optical axis of O2. The position of O3 is adjusted to produce an unclipped, collimated output beam, with collimation verified using a shear interferometer. (D) Placement and alignment of the 200 mm tertiary tube lens (TL3, TTL200-A), with a threaded iris (SM2D25D) used to ensure the beam passes centrally through the lens. (E) Redirecting the emission from TL3 using mirror M13 to guide the beam toward the camera while avoiding any clipping. A neutral density (ND) filter wheel is placed before the camera to prevent sensor damage. The camera is mounted at the focal plane of TL3 and aligned to ensure a focused laser spot at the centre of the field of view.

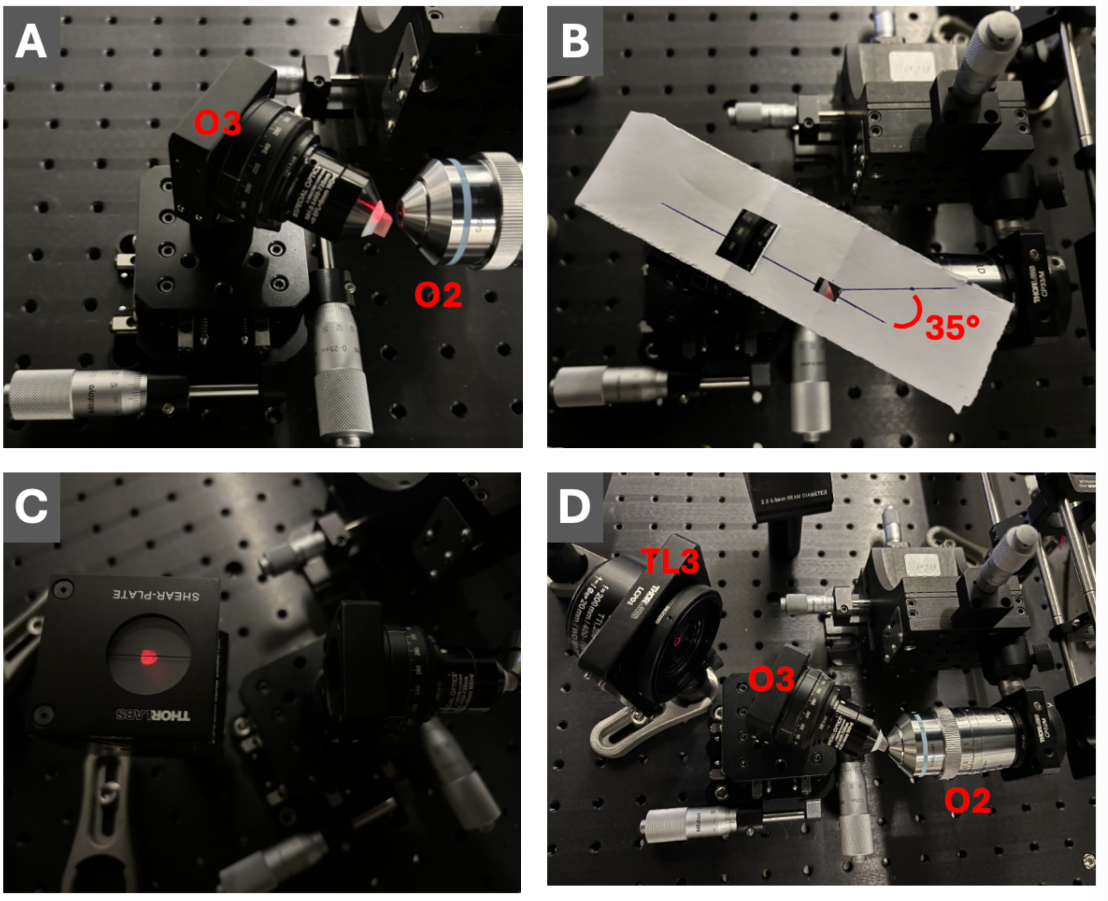


**Photo 14: Alignment of the emission path in the tilted configuration. (Step 124-125)**
(A,B) Mounting O3 at a 35-degree tilt relative to the optical axis of O2. A paper template is used to guide the rotation angle. Adjust O3's position and orientation to achieve an unclipped, collimated output beam. (C) Verifying collimation of the output beam from O3 using a shear interferometer. (D) Positioning the tertiary tube lens (TL3) after O3 to direct the emission path toward the camera, ensuring the output beam remains well-collimated.
