## Supplementary Information for "Constructing a single-objective oblique plane microscope (OPM) for fast, multi-colour, high-resolution volumetric fluorescence imaging"

### The theoretical NA calculation

The theoretical NA of the OPM system is primarily determined by the collection angles of the secondary (O2) and tertiary (O3) objectives. The half-opening angle of the secondary 40× 0.95 NA air objective is 71.8°, while the tertiary 1.0 NA objective has a half-opening angle of 41.5°, matching the critical angle for total internal reflection at an air-glass interface:

$$\theta_c = sin^{-1}\frac{n_{O2}}{n_{O3}} = sin^{-1}\frac{1}{1.51} = 41.5° \quad (1.21)$$

This indicates that O3 can theoretically collect the full emission cone from O2.

**Figure S1** illustrates four situations involving varying degrees of tilt in O3. When O2 and O3 are aligned without tilt, marginal rays (red and blue) are refracted toward the optical axis without clipping, allowing the system to reach the full theoretical collection NA. In this scenario, the half-opening angle at the glass tip of O3 reduces slightly to 39.2°, but this does not limit the effective collection.

Introducing a tilt between O2 and O3, as is common in OPM's remote-refocusing design, affects the collection efficiency. At an 18.2° tilt, the blue marginal ray reaches the air-glass interface at the critical angle, fully entering O3. This angle represents the maximum tilt before the emission cone from O2 begins to be clipped.

In the employed OPM configuration with a 35° tilt, clipping occurs at the air-glass interface, reducing the blue marginal ray's half-opening angle to 55°, while the red marginal ray maintains the full 71.8° angle. Consequently, the angular range along the tilt direction becomes 55° + 71.8° = 126.8°, resulting in an effective NA of 1.25, slightly lower than the nominal 1.35 NA. Perpendicular to the tilt direction (along the Y-axis, Figure S1), no clipping occurs, preserving an effective NA of 1.33.

In an extreme case with a 45° tilt, the angular range further decreases to 45° + 71.8° = 116.8°, reducing the effective NA to 1.19. Although larger tilts can enhance imaging depth, they compromise the effective NA and light collection efficiency.

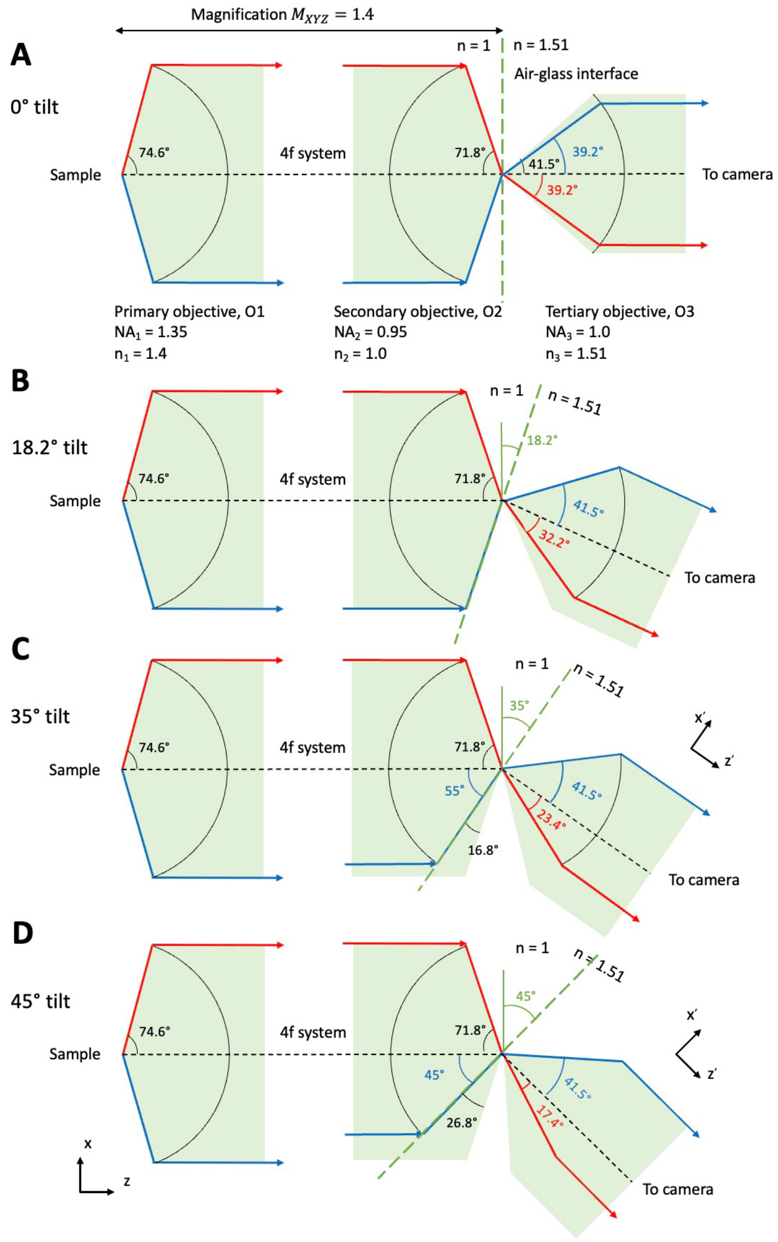

**Figure S1:** Theoretical NA characterisation involving varying degrees of tilt in the Snouty objective. (A) At the 0-degree tilt. (B) At the 18.2-degree tilt. (C) At the 35-degree tilt. (D) At the 45-degree tilt.

### DAQ box connection and LabVIEW control

The galvo mirror (GVS201, Thorlabs) and image acquisition are controlled using a custom LabVIEW program. A cost-effective DAQ box (U3-LV, LabJack) is employed for signal generation and acquisition. The connection is provided in **Figure S2**.

Two analogue control ports on the galvo mirror controller are connected to the DAC0 and DAC1 outputs of the DAQ box. Additionally, the ground of the galvo mirror controller must be connected to the GND terminal of the DAQ box to complete the circuit.

For synchronised galvo scanning and image acquisition, the camera is operated in Edge Trigger Mode, where the rising edge of an external trigger initiates frame acquisition. This trigger signal is generated via the FIO4 port of the DAQ box. The camera's TRIG RDY output is connected to the FIO5 port, allowing the system to monitor when the camera is ready to receive the next trigger. Once the LabVIEW program detects a high signal from FIO5, it generates a rising edge from FIO4 to initiate the next frame.

The custom LabVIEW program also includes functions for automatically scanning the galvo mirror and manually setting it to specific angles, facilitating system alignment and testing. The mirror scanning interval and scanning range can be adjusted by modifying the Step Size, Central Voltage, and the number of Extra Scan (**Figure S3**).

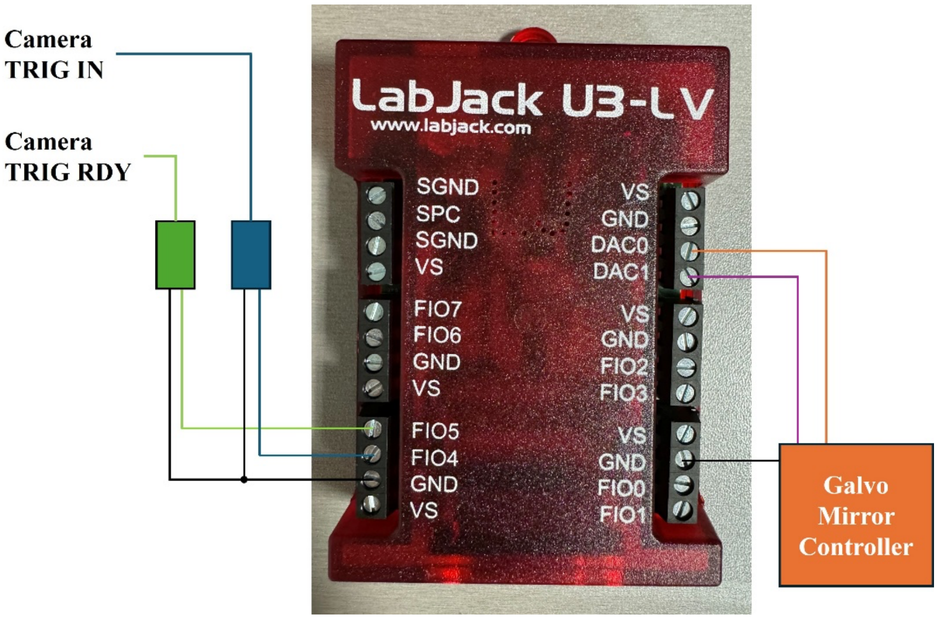

**Figure S2:** Connection diagram for synchronised galvo mirror control and image acquisition using the LabJack U3-LV DAQ. DAC0 and DAC1 output analogue control signals to the galvo mirror controller, while FIO4 and FIO5 are used for camera triggering. FIO4 sends a trigger signal to the camera's TRIG IN port, and FIO5 receives the camera's TRIG RDY signal to indicate readiness for the next frame. Ground (GND) connections are shared to complete the circuit. The green and blue rectangles represent BNC-to-terminal block adaptors used to interface with the camera's TRIG IN and TRIG RDY lines.

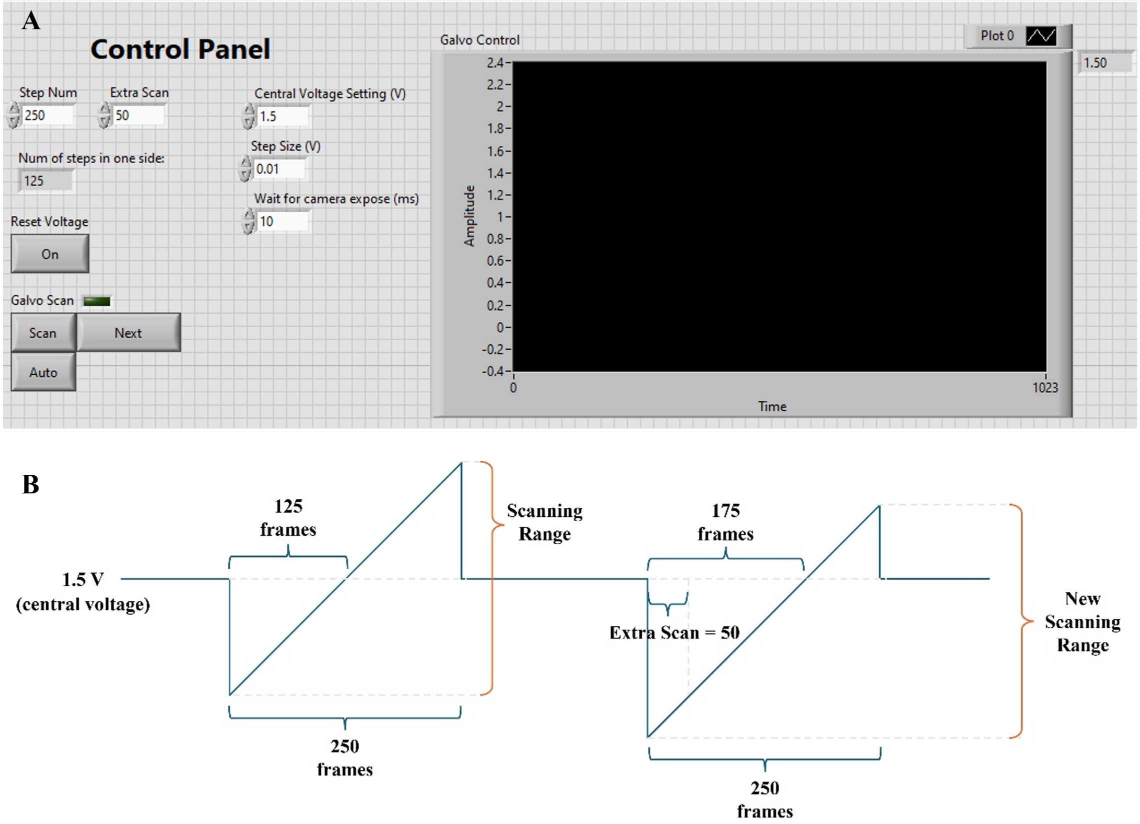

**Figure S3:** Galvo scanning control interface and scan voltage profile. (A) User interface of the custom LabVIEW program used for controlling galvo mirror scanning. Parameters such as step number, extra scan count, central voltage, step size, and camera exposure delay can be adjusted by the user. Real-time voltage output is plotted in the "Galvo Control" window during scanning. (B) Schematic representation of the galvo scanning voltage over time. The scan consists of 250 steps, with 125 frames acquired per direction around a central voltage (1.5 V in this example). An extra scan value of 50 shifts the scanning centre, resulting in a new scanning range. This adjustment allows for fine control over the scanned imaging volume.

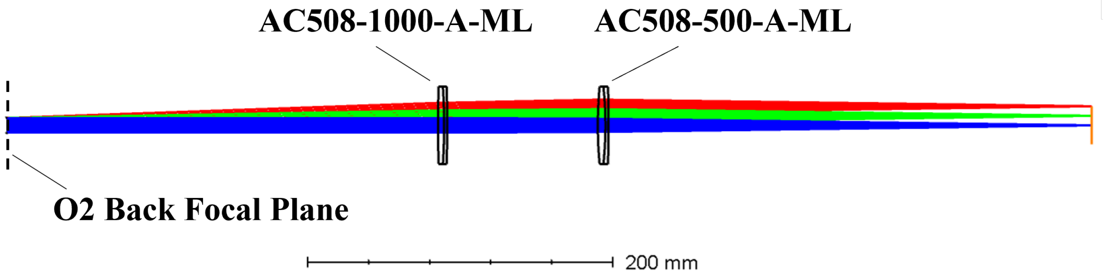

**Figure S4:** Optical layout of the TL2. The TL2 configuration consists of two achromatic doublets (AC508-1000-A-ML and AC508-500-A-ML, Thorlabs) that relay the beam from the back focal plane of objective O2. Ray traces are shown for incident angles of 0°, 1°, and 2°, with ray colour corresponding to different field positions. The 200 mm scale bar indicates the physical length of the optical path for reference.

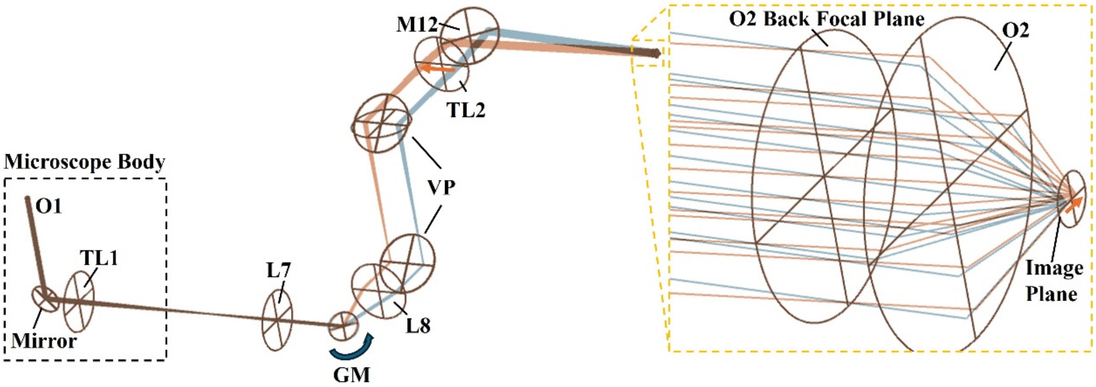

**Figure S5:** Due to the conjugate pupil relay alignment, the beam position at the back focal plane of O2 remains stationary during galvo mirror scanning. As a result, the beam is refocused by objective O2 at its front focal plane (the image plane), causing the focal spot to sweep laterally across the focal plane of O3 (not shown in this figure).

### Fluorescent microsphere in agarose preparation

TetraSpeckTM microspheres, fluorescent blue/green/orange/dark red (T7279, ThermoFisher), with a diameter of 0.1 µm, were embedded in a 3% agarose gel (A9414-5G, Sigma) in PBS on a No.1.5 glass coverslip at a concentration of 1:200. The sample was excited sequentially with lasers at wavelengths of 638 nm and 488 nm, with the corresponding emission filters adjusted accordingly. A total of 150 light-sheet scans were acquired for each excitation wavelength. The raw data was deskewed with home-written Python code without any additional post-processing steps. The FWHM in 3D was calculated for beads located in different areas of the FOV using a 1D Gaussian fit along each axis. The final FWHM values were obtained by averaging the measurements of ten beads selected from the central FOV, where the axial resolution was consistent.

### Cardiomyocyte sample preparation

Adult rat ventricular cardiomyocytes (ARVMs) were isolated from male Sprague–Dawley (SD) rats (150-250 g) using a Langendorff perfusion system. Following isolation, approximately 10,000 cardiomyocytes were seeded onto 35 mm glass-bottom dishes (P35G-1.5-14-C, MatTek) pre-coated with laminin (Cat.#3446-005-01, Cultrex).
Cells were initially incubated for 1 hour at 37 °C in a humidified atmosphere with 5% $CO_2$ in Minimum Essential Medium (MEM; Cat.#31095029, Gibco™) supplemented with 10% fetal bovine serum (FBS; Thermo Fisher) and 1% Antibiotic-Antimycotic Solution (Cat.#A5955, Sigma). After incubation, the medium was replaced with serum-free MEM for subsequent treatments.
To label membrane structures and the actin cytoskeleton, cells were stained with Wheat Germ Agglutinin (WGA) Alexa Fluor™ 488 conjugate (Cat.#W11261, Invitrogen™) and CellMask™ Deep Red Actin Tracking Stain (Cat.#A57245, Invitrogen™), respectively. WGA-488 was used at a final concentration of 10 µg/mL in high potassium buffer (120 mM K-gluconate, 25 mM KCl, 2 mM $MgCl_2$, 1 mM $CaCl_2$, 2 mM EGTA, 10 mM HEPES, 10 mM glucose, adjusted to pH 7.4). CellMask™ Deep Red was first reconstituted in anhydrous dimethyl sulfoxide (DMSO) to make a 1 mM stock solution and subsequently diluted 1:1000 in high potassium buffer to prepare a 1 µM

working solution.
Cells were washed twice with high potassium buffer, then incubated with the combined staining solution for 30 minutes at room temperature in the dark. Following staining, cells were rinsed twice with the high potassium buffer and immediately imaged.

### Diatom sample preparation

The marine diatom Coscinodiscus radiatus (CCMP312) was obtained from the Provasoli–Guillard National Centre for Marine Algae and Microbiota (NCMA, USA). Cultures were grown in artificial seawater medium under controlled environmental conditions at 23 °C, with a 12 h light / 12 h dark cycle and an illumination intensity of 50 $\mu mol \cdot m^{-2} \cdot s^{-1}$, at the Silwood Park campus of Imperial College London. Artificial seawater was prepared using acid-cleaned laboratory materials under aseptic conditions, using American Chemical Society (ACS)-grade or higher purity reagents. The medium was 0.2 µm-filtered using polycarbonate filters (Merck Millipore Ltd.) and stored at 4 °C in the dark. Cultures were maintained by transferring 5 mL of existing culture into 35 mL of fresh medium in new 40 mL culture flasks every two weeks under sterile conditions in a microbiological safety cabinet.
For staining, diatom cells were first incubated in an Eppendorf tube with a 100 µM working solution of SYTO® 9 green-fluorescent nucleic acid stain (Cat.# S34854, Invitrogen™) for 15 minutes at room temperature in the dark. After the incubation, the supernatant was carefully removed, and the pellet was resuspended in 1 mL of fresh culture medium. The diatoms were allowed to settle down for 15 min and the wash process was repeated three times. Then 800 µL of supernatant was removed, the pellet was mixed in the remaining 200 µL of medium and applied to 35 mm glass-bottom dish (P35G-1.5-14-C, MatTek) prepared as follows.
Dishes were first coated with 200 µL of poly-L-lysine solution (P4707, Sigma), incubated for 15 minutes, the excess removed, and then coated with laminin (Cat.#3446-005-01, Cultrex) to promote cell adhesion. The applied diatoms were let 15 min to adhere and transferred to the microscope where 2 mL of culture medium was added carefully not to disturb the cells. Diatoms were immediately imaged after preparation.

### User guide for the galvo mirror control

**1. Hardware setup**
Connect the DAQ box (U3-LV, LabJack) to the camera and galvo mirror controller according to the wiring instructions provided in this paper (see Figure S2).

**2. Configure parameters in the LabVIEW program before running**
Set all relevant values in the control panel. Below is a description of key parameters:
Step Num: Determines the total number of steps for one full galvo mirror scanning volume. Typical value: 252.
Step Size: Determines the voltage interval between each step during galvo mirror scanning. Typical value: 0.01.
Extra Scan: Shifts the centre of the scanning range laterally (see Figure S3).
Wait for camera exposure: Set this equal to the camera exposure time (in milliseconds).
Central Voltage Setting: Defines the central voltage for scanning and also the voltage applied when the "Reset Voltage" button is pressed. Typical value: 1.5V.

**3. Run the LabVIEW program**
Once all parameters are set, run the LabVIEW program to begin control of the galvo mirror.

**4. Galvo mirror control during runtime**
Reset Voltage: Pressing this button sets the galvo mirror voltage to the "Central Voltage Setting" at any point during program execution. Ensure this is turned off during image acquisition.
Scan Modes:
Scan → Auto → Next: Enables continuous scanning across the full voltage range.
Scan → Next: Enables manual step-by-step scanning; click "Next" to advance each step.

**5. During image acquisition**
Ensure that both the "Reset Voltage" and all Scan controls are turned off to ensure correct galvo movements during imaging.

### Micro-Manager 2.0 for 3D acquisition

Micro-Manager is an open-source software for the control and automation of microscope hardware. It works with microscopes from all four major manufacturers (Leica, Nikon, Olympus, and Zeiss), most scientific-grade cameras and many peripherals used in microscope imaging. Download available at: [https://micro-manager.org/Download_Micro-Manager_Latest_Release](https://micro-manager.org/Download_Micro-Manager_Latest_Release)

#### Step 1: Set up the configuration

1. Open Micro-Manager 2.0, select the MMConfig_demo.cfg configuration file.
2. Go to Device > Hardware Configuration Wizard… and select **Create new configuration.**

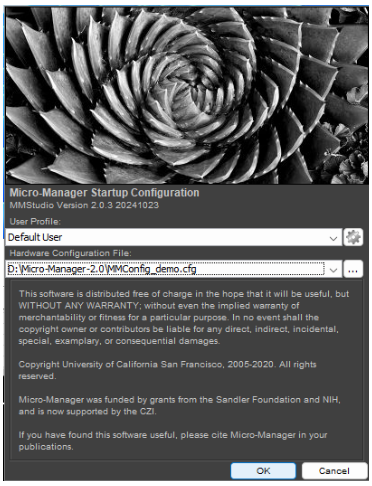

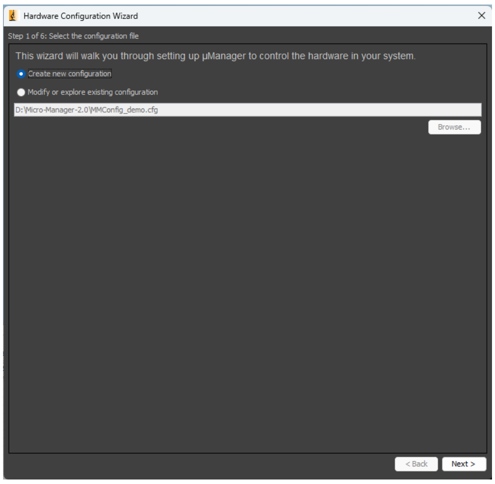

3. Make sure the camera, shutters and the xy stage are all switched on. Select the devices to add to the configuration and click Add…

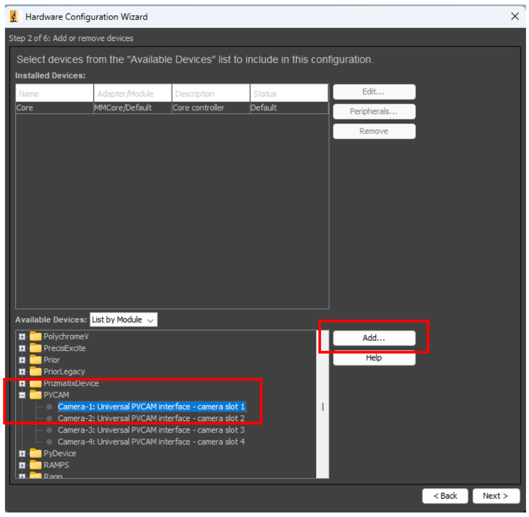

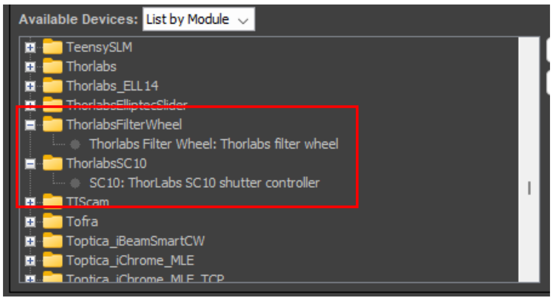

4. Once all devices have been added to the list, click **Next** and follow the prompts to complete the setup. Save the configuration file to your local drive.

**Step 2: Acquire a volumetric image stack**

1. Select the position of interest using the **Live** function in Micro-Manager 2.0. Set the **Exposure** time as desired and set the **Shutter** to **Auto**.

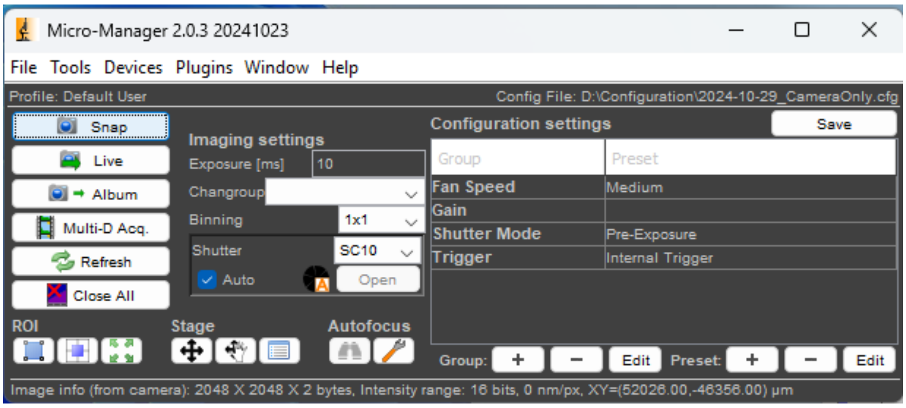

2. Stop Live to avoid photobleaching to the sample.
3. Open the LabVIEW control panel. Set the **Step Num** to your desired number of scan steps (e.g., 250 steps ≈ 145 μm with 2×2 binning). Ensure the camera exposure matches the **Exposure** time set in Micro-Manager.

**Note:** The scan centre can be adjusted by changing the **Central Voltage Setting**, but this is generally not recommended.

4. In Micro-Manager, click **Multi-D Acq**. Set the **Time Points** to **Step Num** in LabVIEW (e.g., for 250 scan steps, set the **Time Points** to 250).

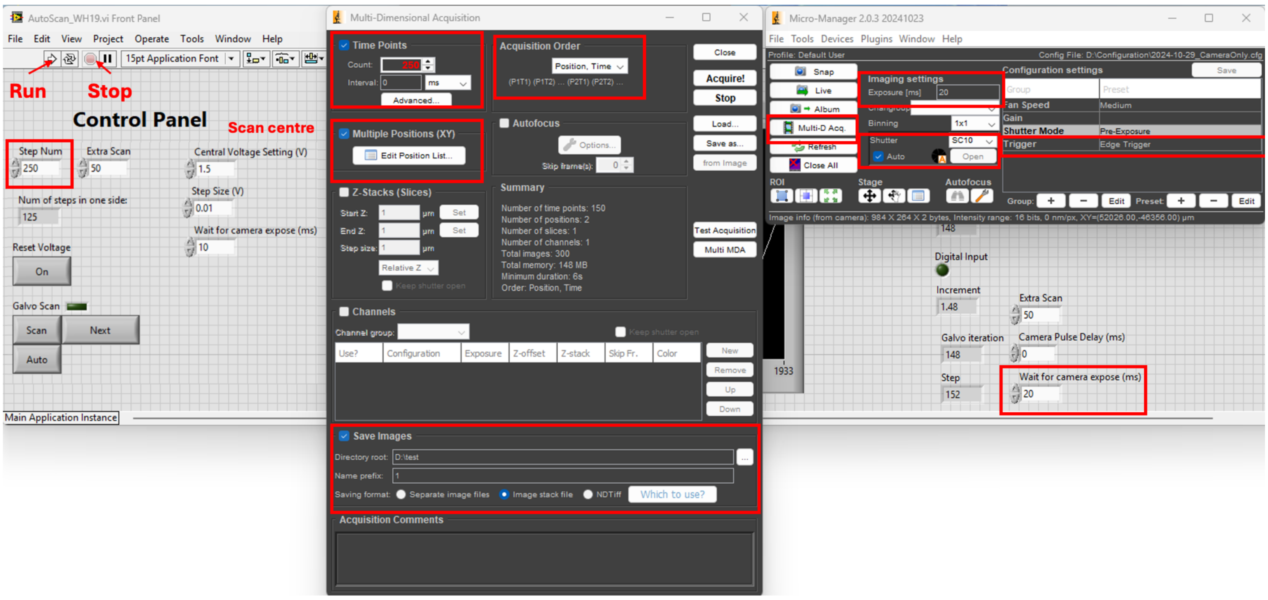

5. You can use the **Multiple Positions (XY)** option for automated scanning of several locations, and specify the directory for saving image stacks.
6. To begin acquisition, set **Camera-1-TriggerMode** (found in **Device > Device Property Browser...**) to **Edge Trigger**. Then, click **Run** in the LabVIEW program, and click **Acquire!** in the Micro-Manager Multi-Dimensional Acquisition window.

Note: To use **Live** function in Micro-Manager, set the trigger mode back to **Internal Trigger**. For volumetric acquisition, always set the trigger mode to **Edge Trigger**.

#### 3D reconstruction of an acquired volumetric stack

As illustrated in Table 2, the imaging volume is at an angle with the sample and therefore the raw image stacks captured by the camera must undergo deskewing to enable accurate 3D reconstruction of the target volume. The principle behind deskewing has been previously described, see Table 2. In brief, two key parameters are required for this process: the tilt angle between O2 and O3 and the pixel step size between adjacent frames relative to the camera pixel size. Using these parameters, each image stack can be computationally reconstructed into a 3D volume with isotropic voxel dimensions along the x, y, and z axes.

The tilt angle of our designed OPM system is 35°. The **pixel step size** can be determined as follows:

1. Acquire a volumetric stack of 0.1 µm fluorescent microspheres embedded in agarose.
2. Use a localisation detection algorithm to determine the pixel coordinates of the microspheres in each frame. For example, here we use the **Peak Fit** function in the **GDSC SMLM** plugin in Fiji to extract the coordinates (Figure S6A).
3. Zoom in to the central region of the Peak Fit result (Figure S6B) and identify a track of fitted dots (Figure S6C). This track corresponds to the position of the same microsphere in successive frames during a volumetric scan.
4. Record the Y-coordinates of the first and last dots and calculate the average separation.
5. The pixel step size to use in the analysis script can then be calculated as:

$$Pixel\ step\ size = S\cos 35° = \frac{(y_2 - y_2)}{No.of\ gaps}\cos 35°$$

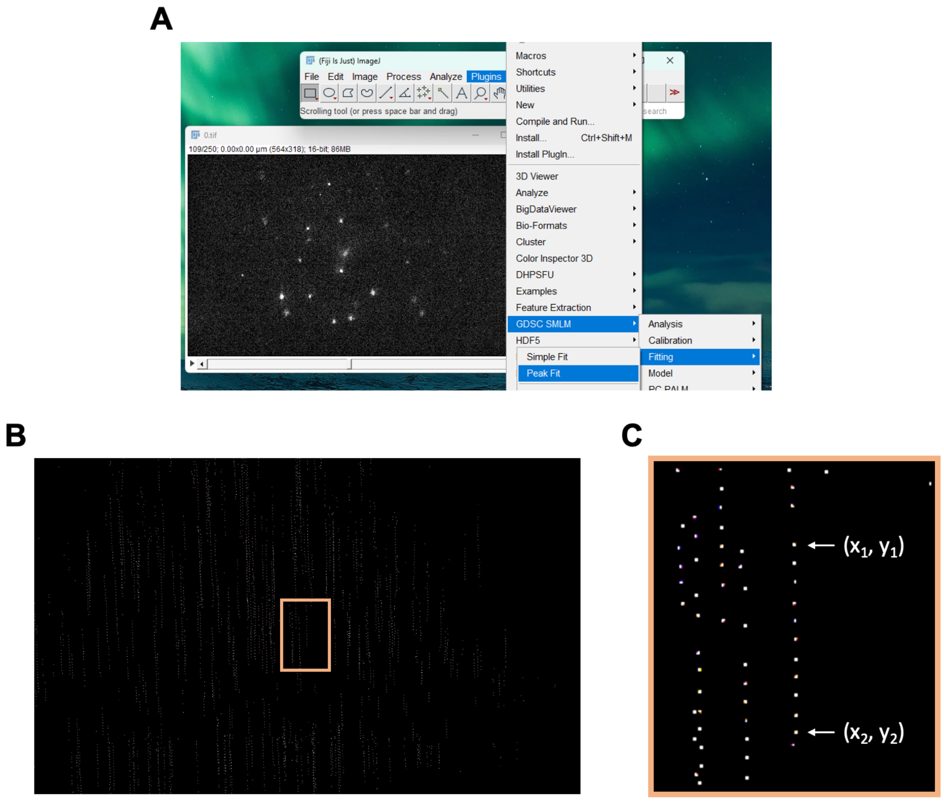

**Figure S6: Procedure to find the pixel step size for the analysis script.** (A) The Peak Fit function in the GDSC SMLM plugin in Fiji was used to extract the coordinates of fluorescent microspheres acquired in a volumetric stack. (B) Output of the Peak Fit analysis, where each bright dot corresponds to a detected microsphere. (C) Enlarged view of the region of interest highlighted in (B), showing a track of fitted dots used for pixel step size calculation.

Beyond deskewing, additional image processing steps, such as cropping, background subtraction, reslicing, and 3D projection, are performed to enhance visualisation and facilitate downstream analysis.

Here, we provide an example Python script for automated processing of multiple image stacks acquired during an experiment. It scans a root directory containing subfolders for different stage positions, each with TIFF image stacks, and creates a new analysis directory that mirrors the original structure. The script then processes each image stack by cropping to a user-defined region of interest using Fiji, subtracting background via the rolling ball algorithm, deskewing based on the specified light-sheet tilt angle and inter-frame spacing to generate isotropic 3D volumes, reslicing the reconstructed stack along the z-axis, and producing 3D projection images rotating about the y axis with a fixed angle increment. In particular, multiprocessing is used to accelerate the deskewing step, and all stages are saved in separate folders to enable viewing and analysis. The analysis script can be downloaded from: https://github.com/Zui409/Complete-suite-for-single-objective-OPM
